# Prefrontal working memory signal controls phase-coded information within extrastriate cortex

**DOI:** 10.1101/2024.08.28.610140

**Authors:** Mohsen Parto-Dezfouli, Isabel Vanegas, Mohammad Zarei, William H. Nesse, Kelsey L. Clark, Behrad Noudoost

**Affiliations:** Max Planck Institute for Biological Cybernetics, 72076 Tübingen, Germany; Ernst Strüngmann Institute (ESI) for Neuroscience in Cooperation with Max Planck Society, 60528 Frankfurt, Germany; Department of Ophthalmology and Visual Sciences, John Moran Eye Center, University of Utah, Salt Lake City, UT, United States; Biosciences Institute, Newcastle University, Newcastle upon Tyne, UK

**Keywords:** neural phase code, working memory, top-down control, prefrontal cortex, neural oscillations

## Abstract

In order to understand how prefrontal cortex provides the benefits of working memory (WM) for visual processing we examined the influence of WM on the representation of visual signals in V4 neurons in two macaque monkeys. We found that WM induces strong β oscillations in V4 and that the timing of action potentials relative to this oscillation reflects sensory information-i.e., a phase coding of visual information. Pharmacologically inactivating the Frontal Eye Field part of prefrontal cortex, we confirmed the necessity of prefrontal signals for the WM-driven boost in phase coding of visual information. Indeed, changes in the average firing rate of V4 neurons were correlated with WM-induced oscillatory changes. We present a network model to describe how WM signals can recruit sensory areas by inducing oscillations within these areas and discuss the implications of these findings for a sensory recruitment theory of WM through coherence.

## Introduction

Our capacity to dynamically interact with the world around us based on our own needs, priorities and goals is an example of our cognitive flexibility. The goals and plans preserved by working memory (WM) are capable of altering our perceptions of and actions toward the world around us. Thus, understanding how our plans alter the representation of sensory information can reveal the neural basis of cognitive flexibility. Prefrontal cortex (PFC) is believed to be one of the sources controlling sensory signals according to goals and plans^1-7^. In this study, we are specifically examining how WM information sent from PFC can influence the representation of sensory information in visual areas in macaque monkeys. We have already shown that the Frontal Eye Field (FEF) part of PFC sends a direct WM signal to extrastriate area V4^8^, and visual areas manifest the WM content in their oscillatory behavior^9^. Within the FEF, WM alters the efficacy of V4 inputs as well, and these inputs mostly target neurons believed to be involved in the transformation of visual information into motor action^10^. The finding that extrastriate visual areas receive the content of WM, and that information sent from these areas to prefrontal cortex is undergoing a visuomotor transformation, provide a clear picture of the sequence of events giving rise to the benefits of WM. However, exactly how this top-down WM signal enhances the representation in visual areas is not known, considering that neurons in extrastriate visual areas show little or no change in their firing rate in response to WM content^11-15^. Thus, how WM impacts visual representations is crucial for understanding the neural code, since the behavioral consequences of WM^16-22^ suggest that the representation altered by WM content is likely to be the representation that our behavior relies on.

Knowing that the FEF sends a spatially-specific signal to V4 carrying the content of WM^8^, we studied the responses of V4 neurons while top-down WM is directed to their receptive field (RF) or elsewhere, under the conditions in which FEF activity is intact or disrupted using pharmacological manipulation. V4 neurons were provided with a bottom-up visual input with the goal of understanding how a top-down WM signal can alter the representation of bottom-up information. This arrangement revealed that WM primarily enhances the phase coding of visual information in V4. Pharmacological inactivation demonstrated that FEF activity is necessary for this phase-dependent representational enhancement in V4. We also show that average firing rate modulations within visual areas correlate with WM-induced oscillations within these areas, suggesting a possible role of WM-dependent oscillations in generating the other signatures of WM within sensory areas. These results support a WM model in which sensory areas can potentially be recruited by higher areas without needing to change the average firing rate of sensory neurons.

## Results

### WM modulates oscillatory power and spike timing in V4, but not average firing rates

In order to assess the impact of WM on the visual representation, we recorded local field potentials (LFPs) and neuronal activity in extrastriate area V4 during a spatial WM task with visual signals presented to neurons independent of the WM demand (see Methods; Fig. 1A). The animal had to remember a visual cue presented either inside the V4 RF or in the opposite hemifield (memory IN and OUT conditions, respectively), and following a delay period, made a saccade to the remembered location to receive a reward. The background stimulus could have one of four orientations and three levels of contrasts (or no contrast, the classic memory-guided saccade (MGS) task), allowing us to examine the interaction between bottom-up sensory information and top-down WM signals. We recorded 145 V4 neurons across 88 recording sessions; most of our analysis focuses on their responses during the last 700ms of the delay period of the WM task.

**Figure 1.**
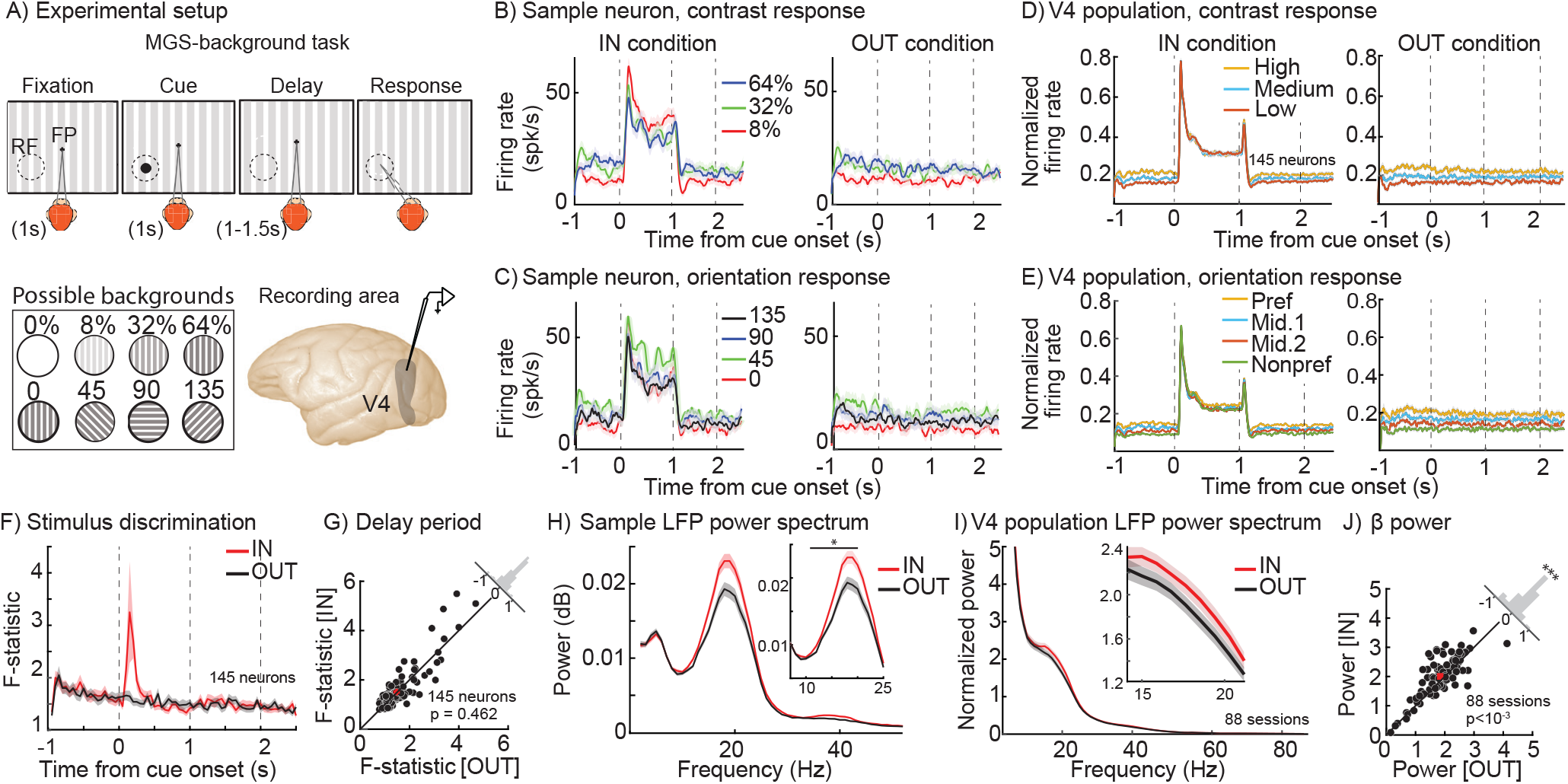
WM alters β oscillatory power but not firing rates in V4. A) Memory-guided saccade with background (MGS-background) task. The monkey fixated and a peripheral visual cue appeared (Cue). The monkey maintained fixation while remembering the cue location (∼1s; Delay), and after the fixation point disappeared, executed a saccadic eye movement to the remembered location (Response) to receive a reward. Throughout the task, there was a task-irrelevant, full-field oriented bar background; the background contrast ranged from 0-64%, in one of 4 orientations. The memory location was either inside the extrastriate RF (IN condition, shown) or 180 degrees away (OUT condition). Neurophysiological recordings of spiking and LFP activity were made from extrastriate visual area V4, with linear array or single electrodes. B-C) Mean firing rate of a sample neuron over time for three different contrasts (B) or orientations (C) of the background stimulus, for the IN (left) and OUT (right) conditions. Shaded areas in all panels show standard error of mean (SEM). D-E) Mean firing rate of the population of 145 neurons over time for three different contrasts (low, medium, and high contrast, D) and four different orientations (preferred, nonpreferred, and middle 1 & 2 orientations, E) for the IN condition (left) and OUT (right) conditions. F) Time course of mean F-statistic values across 145 neurons, based on a one-way ANOVA for discrimination between 12 stimulus conditions for the IN (red) and OUT (black) conditions. G) Scatter plot of F-statistic averaged in the last 700ms of the delay period for each session, for the IN vs. OUT conditions. Histogram in the upper right shows the distribution of change in F-statistic (OUT-IN) across sessions. H) Mean power spectrum of the LFP recorded from the same channel as the sample V4 neuron in (B), during the delay period for the IN (red) vs. OUT (black) conditions. Inset panel shows 8-25 Hz. Asterisk indicates a significant difference (p<0.05) in the range shown. I) Mean power spectrum of population of the V4 LFPs (88 sessions) during the delay period for the IN (red) vs. OUT (black) conditions. Inset shows the power spectrum between 14-22 Hz. J) Scatter plot of power spectrum averaged in the β range for each session, for the IN vs. OUT conditions. Histogram in the upper right shows the distribution of change in power (OUT-IN) across sessions (***, p <0.001).

As expected, V4 neurons were capable of signaling the properties of the background stimulus present in their RF. The left panel in figure 1B shows the response of a sample V4 neuron during the IN condition. During the fixation period and the cue period, the neuron exhibited sensitivity to different contrast levels of the background stimulus projected to its RF (main effect of contrast: F_Fixation_=41.744, p<10^−16^; F_Cue_=14.541, p<10^−6^; one-way ANOVAs). The neuron’s contrast sensitivity was still manifested in its firing rate during the delay period (F_IN_=19.950, p<10^−8^; One-way). As shown on the right side of figure 1B, this neuron also showed contrast sensitivity during the delay period of the OUT condition (F_OUT_= 17.851, p<10^−7^; One-way ANOVA). Importantly, the contrast sensitivity during the delay period was not significantly different between the IN vs. OUT conditions (F_condition_= 5×10^−4^, p=0.981; F_contrast_=37.540, p<10^−16^; F_interaction_=0.301, p=0.742; Two-way ANOVA). Similarly, the neuron was selective for the orientation of the background stimulus. The neuron exhibited orientation sensitivity during the fixation period as well as the delay period in both IN and OUT conditions (main effect of orientation: F_Fixation_=16.401, p<10^−8^; F_IN_=10.410, p<10^−5^; F_OUT_=19.014, p<10^−10^; One-way ANOVAs; Fig. 1C). As with contrast, the content of WM (IN vs. OUT condition) did not significantly change the neuron’s orientation sensitivity during the delay period (F_Condition_= 5×10^−4^, p=0.982; F_Orientation_=28.082, p<10^−16^; F_Interaction_=0.810, p=0.492; Two-way ANOVA). Thus, this sample neuron exhibited contrast and orientation sensitivity for the background stimulus; however, WM did not alter the stimulus information reflected in the neuron’s firing rate.

The same pattern was observed in the population of 145 V4 neurons: neurons’ firing rates reflected the properties of the background stimulus, but this capacity was not affected by WM. The response to different background stimuli was similar between the fixation period, delay IN, and delay OUT conditions, both for contrast (F_Contrast_ =16.923, p<10^−7^; F_Condition_ =0.311, p=0.733; F_Interaction_ =0.400, p=0.811; Two-way ANOVA) and orientation (F_Orientation_ =23.294, p<10^−14^; F_Condition_ =0.770, p=0.465; F_Interaction_ =0.202, p=0.976; Two-way ANOVA) (Fig. 1D, 1E; see also Fig. S1A-B). Figure 1F shows the ability of V4 neurons to discriminate between all 12 stimuli of various contrasts and orientations across time. Overall, the ability of V4 neurons to discriminate between various background stimuli was not altered during the delay period of the IN vs. OUT conditions (discriminability_IN_=1.514±0.783, discriminability_OUT_=1.482±0.680, p=0.462; Fig. 1G; see also Fig. S1C-D). Therefore, whereas sensory information is reflected in the firing rate of V4 neurons, the impact of spatial WM on the sensory representation is not traceable by this neural signature.

In contrast to the lack of WM-driven change in firing rates, we found that V4 LFP oscillations strongly reflect the content of WM. WM-driven changes could be seen at the level of the average LFP (Fig. S2), and more robustly in the LFP power spectrum. Figure 1H shows the LFP power spectrum during the delay period for the same channel of recording as of the example neuron shown in figure 1B-C. The β (14-22 Hz; see Methods) band power was 0.019±0.013 dB/Hz during the IN condition, which is 18.9% greater than the 0.016±0.012 dB/Hz during the OUT condition (Wilcoxon ranksum, p=0.002). Therefore, although neurons’ firing rates do not change due to WM, the power of the LFP oscillation in the β range reflects the impact of WM in the same recording channel. This phenomenon was observed across the 88 LFP recordings (Fig. 1I): β LFP power during the delay was greater for the IN condition compared to the OUT condition (Power_IN_ =2.001±0.336, Power_OUT_ =1.842±0.333, p<10^−3^; Fig. 1J). Importantly, this WM-dependent enhancement of β power was observed independent of the background stimuli (Fig. S3); the discriminability based on the LFP power spectrum across frequencies is shown for different WM conditions in figure S4. In summary, a visual area known to receive the spatial WM signal exhibits the signature of this top-down signal in its subthreshold LFP activity, but not in neurons’ firing rates.

The LFP reflects a combination of nearby currents, including synaptic inputs^23-26^, and is not directly transmitted along axons to other brain areas in the manner that spikes are. Therefore, we sought to identify whether these WM-dependent oscillatory changes impact any other aspects of spiking activity within V4. Although at the coarse scale of average firing rate there was no change due to WM, at a higher temporal resolution it became evident that the timing of V4 action potentials depended on the phase of WM-induced β oscillations. Figure 2A shows the distribution of spikes generated by a sample V4 neuron across various phases of the β oscillation during the delay period, for memory IN and OUT conditions. The average delay period FR was not different between the two memory conditions (F_Condition_= 0.219, p=0.640; F_Contrast_ =24.777, p<10^−10^; F_Interaction_ =0.180, p=0.835; Two-way ANOVA), but the phase distribution of spikes during the delay period of the IN condition was more concentrated (centered around 175-degree phase) compared to the OUT condition. To quantify the phenomenon, we used the spike-phase locking (SPL) as a measure of how consistently spikes of a neuron are generated at a certain phase. The SPL index varies between 0 (spikes homogenously distributed across phases) to 1 (all spikes occurring at a certain phase). For the sample neuron shown in figure 2A, SPL changed from 0.169 for the OUT to 0.189 for the IN condition. Across the population of 145 V4 neurons, we found a consistent impact of WM on SPL: SPL relative to the β oscillation was significantly greater during the IN condition compared to the OUT condition (n=145 neurons; SPL_IN_ =0.164±0.014, SPL_OUT_ =0.155±0.005, p<10^−3^; Fig. 2B). Conversely, evaluating the consistency of the LFP at the time of V4 spikes, the spike triggered average (STA) LFP also shows this strong coupling of spikes to LFP phase in the β range (Fig. S5). The 0% contrast condition, equivalent to the classic MGS task, also showed significantly greater β power (n = 88 sessions; Power_IN_=1.957±0.0.651, Power_OUT_=1.818±0.733, p=0.006, Wilcoxon signed-rank) and SPL (n=145 neurons, SPL_IN_ =0.135±0.048, SPL_OUT_ =0.119±0.055, p<10^−7^, Wilcoxon signed-rank) for the IN condition (Fig. S6); these findings are similar to a previous report in visual area MT^9^. These results indicate that although WM does not change the average firing rate, it influences V4 spike timing to be more closely aligned with WM-dependent oscillations.

**Figure 2.**
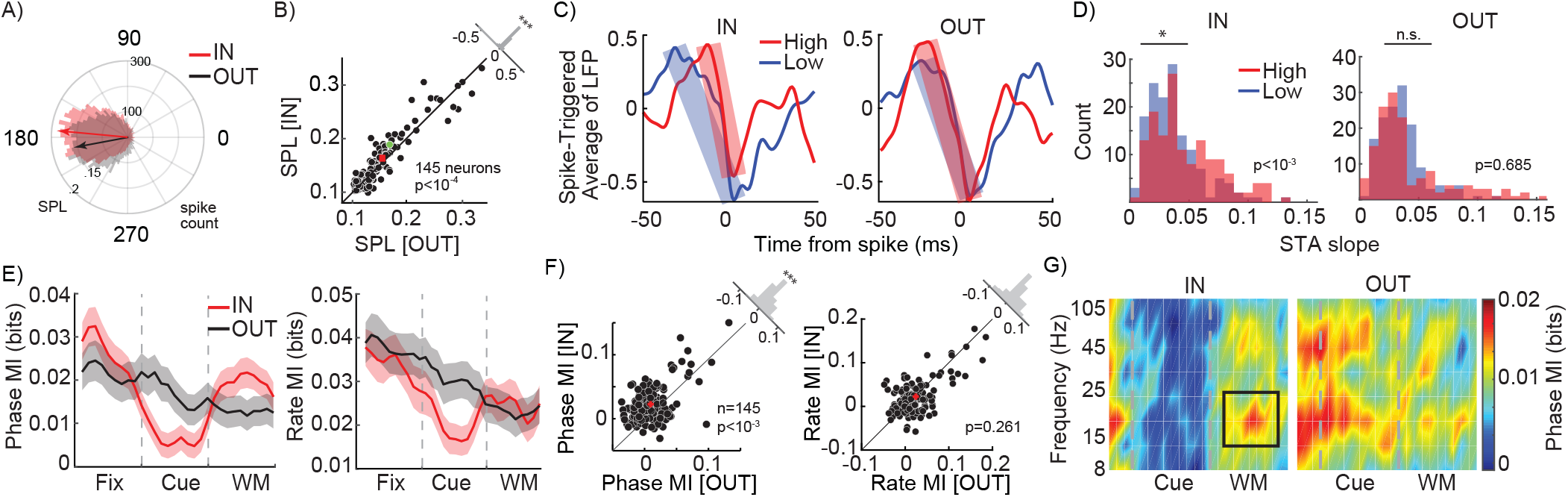
WM alters the sensory representation in extrastriate cortex. A) The distribution of spikes generated by a sample V4 neuron across various phases of β oscillations during the delay period. Arrows show the average of phase distributions for the IN (red) and OUT (black) conditions. B) Scatter plot of SPL in the β range for each neuron, for the IN vs. OUT conditions. Histogram in the upper right shows the distribution of change in SPL (OUT-IN) across neurons. Red cross indicates population mean. Green dot shows the selected sample neuron in A. C) Spike-triggered average of the normalized LFP of a sample neuron during the delay period, for the high contrast (red) and low contrast (blue) background stimuli, for the IN (left) and OUT (right) conditions. Shaded bars indicate the slopes in the falling phase. D) Histogram of the distribution of STA slopes (abs (V_peak_ – V_trough_)/(time_peak_ – time_trough_)) across neurons, for high contrast (red) and low contrast (blue) stimuli, for the IN (left) and OUT (right) conditions. E) Population phase (left) and rate (right) coding over time, for the IN (red) and OUT (black) conditions, based on mutual information (MI) between 12 stimulus conditions. MI was measured in 100ms windows with steps of 100ms. Shaded areas show standard error of mean (SEM). F) Scatter plot of MI using a phase code in the β range (left) and rate code (right) for each neuron, for the IN vs. OUT conditions. Red crosses indicate population mean. Histograms in the upper right show the distribution of differences in MI (IN-OUT) across neurons. G) Phase coding, measured by MI (colorbar), as a function of frequency and time, for memory IN (left) and OUT (right). Black rectangle shows the time and frequency range selected for phase code analysis. (*, p <0.05; **, p <0.01; ***, p <0.001; ns, p>0.05)

### Spatial WM specifically enhances phase coding of visual information

To understand how WM-induced oscillations benefit the sensory representation within V4, we examined whether V4 neurons’ sensitivity was reflected in the timing of their spikes relative to these WM-induced oscillations. As shown in figure 2C, analyzing the average normalized LFP at the time of spiking (the STA) for an example neuron, we observed that in the presence of the high contrast stimulus in the background during the memory IN condition, V4 spikes were associated with a steeper LFP change compared to when a low contrast stimulus was in the background (Slope_High_=0.052, Slope_Low_=0.029). For this neuron, this difference was less prominent during the OUT condition (Slope_High_=0.042, Slope_Low_=0.035). We observed a similar phenomenon across the population. The LFP slope around the time of a spike was significantly sharper when the preferred (high contrast) stimulus was presented in the background, compared to the nonpreferred (low contrast) stimulus, during the IN condition (Slope_High_ =0.047±0.002, Slope_Low_ =0.37±0.003, p<10^−3^; Fig. 2D left). This difference was not observed during the OUT condition (Slope_High_=0.042±0.003, Slope_Low_=0.036±0.003, p=0.686, Fig. 2D right). This indicates that WM-induced oscillations can facilitate the spiking activity in visual areas, consistent with what other groups have shown regarding the role of oscillations as a boost for passing the spiking threshold ^27^. Similar results were observed for preferred versus nonpreferred orientations (Fig. S7).

The idea that a WM-induced oscillation can change the timing of spikes also suggests the possibility that the timing of spikes relative to that oscillation could convey information, referred to as a neural phase code. We used the mutual information (MI) to quantify visual information conveyed by either the phase or rate of spikes to have a side-by-side comparison of a neural phase vs. rate code (see Methods). Figure 2E left shows the average population MI measured based on phase coding across time for both the IN and OUT conditions. The presence of the WM cue reduced both phase and rate coding of background information during the visual period (Phase MI_IN_=0.007±0.001 bits, Phase MI_OUT_=0.014±0.002 bits; p=0.012; Rate MI_IN_=0.019±0.002 bits, Rate MI_OUT_=0.031±0.003 bits; p=0.012). However, maintenance of WM information during the delay period increased the phase coding capacity of the V4 neurons to represent information about the stimulus in their RF, but did not alter their rate coding. Figure 2F shows the capacity of V4 neurons to encode the background stimulus during the IN vs. OUT conditions under phase (left) and rate (right) coding schemes. Phase MI during the delay period of the IN condition was 0.019±0.002 bits, significantly greater than the 0.012±0.002 bits during the OUT condition (p<10^−3^). However, delay period rate MI was not significantly different between the IN and OUT conditions (MI_IN_=0.024±0.003 bits; MI_OUT_=0.022±0.003 bits, p=0.261). We also found that the WM-dependent enhancement of phase-dependent visual representation was limited to the β range oscillations (Fig. 2G) It is imperative to note that despite the seemingly small values of MI (e.g., 0.019 bits), an increase of 54% in phase MI between the IN and OUT conditions means a huge boost in coding capacity due to WM. The MI value can be interpreted as a measure of the rate of statistical learning from incoming data through which sensory decisions can be made. For example, with 0.012 bits per 100 ms phase MI available in the OUT condition, for a population of 100 neurons firing independently it would take 294 ms to fully differentiate 12 stimuli. WM-induced enhancement of phase MI means that the same discriminatory capacity can be achieved within 190 ms with the same number of neurons, or within the same amount of time but with only 65 neurons. Altogether, consistent with the finding that WM mainly modulated oscillations within the β range (Fig. 1I, J), we found that WM mostly improves the phase coding in V4 within the same β range. Thus, WM specifically enhanced β range phase coding in V4, without altering rate coding.

### FEF activity is necessary for the phase-dependent representation within V4 during WM

FEF sends direct projections to V4; these projections arise primarily in the superficial layers of FEF ^28-30^, and are functionally characterized by delay activity reflecting the content of spatial WM^8^. To causally test whether the observed WM-driven phase coding in V4 depended on signals received from the FEF, we recorded from V4 neurons before and after pharmacologically inactivating a portion of the FEF using a small-volume injection of the GABAa-agonist muscimol (Fig. 3A). Localized FEF inactivation is known to impair performance on the MGS task in a spatially-specific manner^31,32^. As shown for an example inactivation session in figure 3B: prior to inactivation, the animal performed well at all locations, and after FEF inactivation performance was disrupted for conditions in which the cue appeared in the left hemifield, contralateral to the inactivated FEF. Figure 3C shows average MGS performance over time at various locations across 33 inactivation sessions: performance for the IN condition and neighboring locations decreased over time following FEF inactivation. For the IN condition, the performance dropped from 90.32±2.62 percent correct before to 68.49±5.39 percent correct three hours after inactivation (p<10^−3^). Across the same 33 sessions, saccade error during the IN condition compared to the OUT condition was not different prior to FEF inactivation (Error_IN_=1.012±0.022, Error_OUT_=0.989±0.023, p=0.520; Fig. 3D), but was significantly greater for the IN condition after FEF inactivation (Error_IN_=1.192±0.042, Error_OUT_=1.033±0.032, p=0.001; Fig. 3D). Similarly, reaction time (RT) increased following FEF inactivation for the IN condition compared to the OUT condition (before inactivation: RT_IN_=1.015±0.003, RT_OUT_ =1.023±0.007, p=0.741; after inactivation: RT_IN_ =1.075±0.015, RT_OUT_ =1.014±0.006, p<10^−4^, Fig. 3E). We found no effect of the background stimulus on the task performance, either in terms of reaction time (Fig. S8A; ANOVA between three contrasts: F_IN_=1.75, p=0.180; ANOVA between four orientations: F_IN_=2.26, p=0.085) or in percent correct (Fig. S8B; ANOVA between three contrasts: F_IN_=0.01, p=0.990; ANOVA between four orientations: F_IN_=0.06, p=0.979).

**Figure 3.**
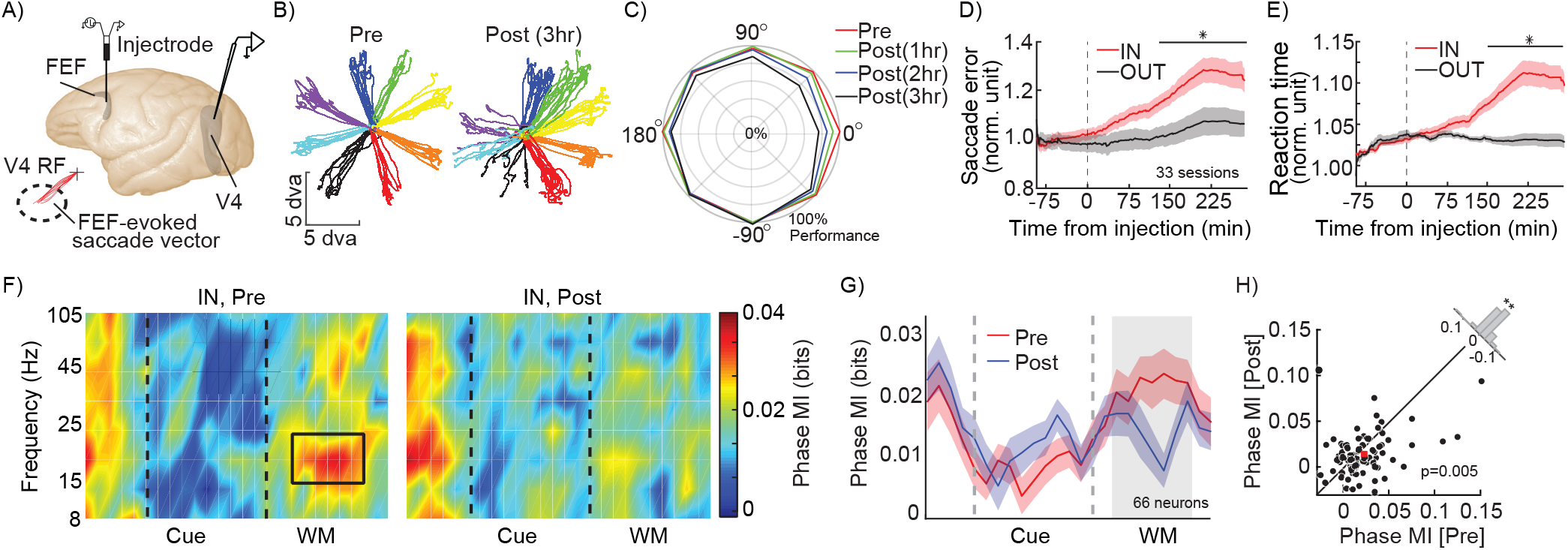
FEF inactivation alters WM behavioral performance and phase coding in visual areas. A) V4 recordings were made before and after infusion of muscimol into FEF. Muscimol injections into FEF were made with a custom microinjectrode, at sites with stimulation-evoked saccade endpoints overlapping with simultaneous V4 recording site RFs. B) Eye traces for 8 MGS target locations, before (left) and after (right) FEF inactivation, for an example session where 0.5 microliter of muscimol was injected into the FEF; performance deficits were localized to the infusion hemifield. C) Average behavioral performance across sessions, at different locations over time following FEF inactivation (red pre-inactivation; green, blue, and black, 1, 2, and 3 hours after inactivation, respectively). Data from each session is aligned so that 0 degrees corresponds to the FEF RF. D-E) Normalized saccade error (D) and reaction times (E) for the memory IN (red) and OUT (black) conditions, over time relative to the FEF inactivation. Black bar indicates times with a significant difference between IN and OUT. Shaded areas show SEM across sessions. F) Heatmap shows phase coding (MI, colorbar) over time and frequency for 66 V4 neurons, for the IN condition, before (left) and after (right) FEF inactivation. Black rectangle indicates time and frequency range considered in (G-H): 14-22Hz, 200-800ms after start of delay period. G) Strength of β phase coding over time, for memory IN, before (red) and after inactivation (blue). The phase-code MI is averaged in the β range. Shading shows SEM across neurons. Gray area indicates time window plotted in (H). H) Scatter plot of β phase MI during the delay period (shaded area in G) of the IN condition for each V4 neuron, before vs. after FEF inactivation. Red square shows population mean. The histogram in the upper right shows the distribution of difference in MI (Pre-Post) across neurons. (*, p <0.05; **, p <0.01; ***, p <0.001; ns, p>0.05)

Next we evaluated the impact of FEF inactivation on V4 activity during WM. We recorded from 66 V4 neurons before and after FEF inactivation. Discriminability of the background stimulus based on V4 firing rates was unaltered, and based on LFP only slightly altered, following FEF inactivation (Fig. S9A-C).

Both the V4 LFP power spectrum and SPL showed a reduction in the β range following FEF inactivation (Fig. S10). In these 66 neurons, prior to inactivation, the impact of WM on phase coding was evident: during the delay period there was significantly stronger phase coding of information in the β range for the memory IN condition (Fig. 2E-G; Phase MI_IN_=0.023±0.004 bits, Phase MI_OUT_=0.018±0.003 bits, p=0.039). Consistent with figure 2E, WM did not alter the strength of rate coding (Rate MI_IN_=0.022±0.005 bits, Rate MI_OUT_=0.020±0.005 bits, p=0.446). Importantly, the phase coded information within the β range during the delay period of the task dropped following FEF inactivation (Fig. 3F). Figure 3G shows the cross section of figure 3F at the β frequency, depicting the dynamics of phase MI over the course of a trial before and after FEF inactivation. The MI values for individual neurons during the delay period of the IN condition for each session are shown in figure 3H; following FEF inactivation, phase-coded MI in the β range dropped from 0.023±0.004 bits to 0.014±0.003 bits (n= 66 neurons, p=0.005; Fig. 3H). The phase coded information within the β range during the delay period of the task does not change following FEF inactivation for the OUT condition (n= 66 neurons, p=0.281; Fig. S11). Rate coding, in contrast, was unaffected by FEF inactivation (IN condition: Rate MI_Pre_=0.022±0.005 bits, Rate MI_Post_0.020±0.004 bits, p=0.880). Thus, WM’s enhancement of phase coding in V4 depended on activity within the FEF.

### Modelling WM-driven changes in phase and rate coding

The finding that WM mainly modulates phase coded information within extrastriate areas fundamentally shifts our understanding of how the top-down influence of prefrontal cortex shapes the neural representation, suggesting that inducing oscillations could be the main way WM recruits sensory areas. However, while this side-by-side comparison of rate and phase coding shows the strength of the latter, several studies have reported an impact of WM on the firing rate of visual neurons^8,15,33-35^. One can argue that a slight increase in firing rate at each stage of visual processing can gradually accumulate to eventually emerge in the form of a robust firing rate change^12^, and that this will be sufficient to support WM. In order to determine the primary means by which WM alters neural representations, we constructed a neural field network model of visual areas during WM. In order to examine the impact of WM on oscillatory and firing rate changes in visual areas, we designed the model to consist of interconnected excitatory and inhibitory units (Fig. 4A, e-cells and i-cells) capable of generating oscillatory activity. To modulate this oscillatory activity, these neural field units received bottom-up and top-down type input. The units were tuned to different stimulus ‘features’ of the bottom-up input (analogous to orientation tuning of V4 neurons in the experimental data, with input strength analogous to contrast). The top-down input was not feature selective, providing a uniform input across the network, with stronger connections to e-cells than i-cells, consistent with what is known about the FEF-V4 circuitry and anatomy^8,36^. A higher strength of WM signal in the model corresponds to the memory IN condition, in comparison to the absence of WM input in the memory OUT equivalent. The model replicates several key features of the experimental data: units reflect sensory information in their phase and rate, WM-enhanced β power, and locking of units’ activity to this oscillation under the influence of WM (Fig. S12). The model’s phase coding of visual stimuli is evident in the relative timing of responses of differently tuned e-cells to an input stimulus (Fig. 4B). Using this model, we can directly compare the magnitude of information encoded by the phase and rate of model units, quantified via information gain (see Methods). We found that not only was information encoded by phase much greater than that encoded by rate, but also that phase and rate information were oppositely affected by changes in WM strength: phase information increased and rate information decreased as the WM signal increased (Fig. 4C). We also found this same divergent pattern between phase and rate codes when measuring coding performance using mutual information (see Fig. S13). Therefore, the quantification of information within a tangible network model revealed that in an oscillating network, a top-down induced oscillation can be detrimental to the rate-dependent representation of information.

**Figure 4.**
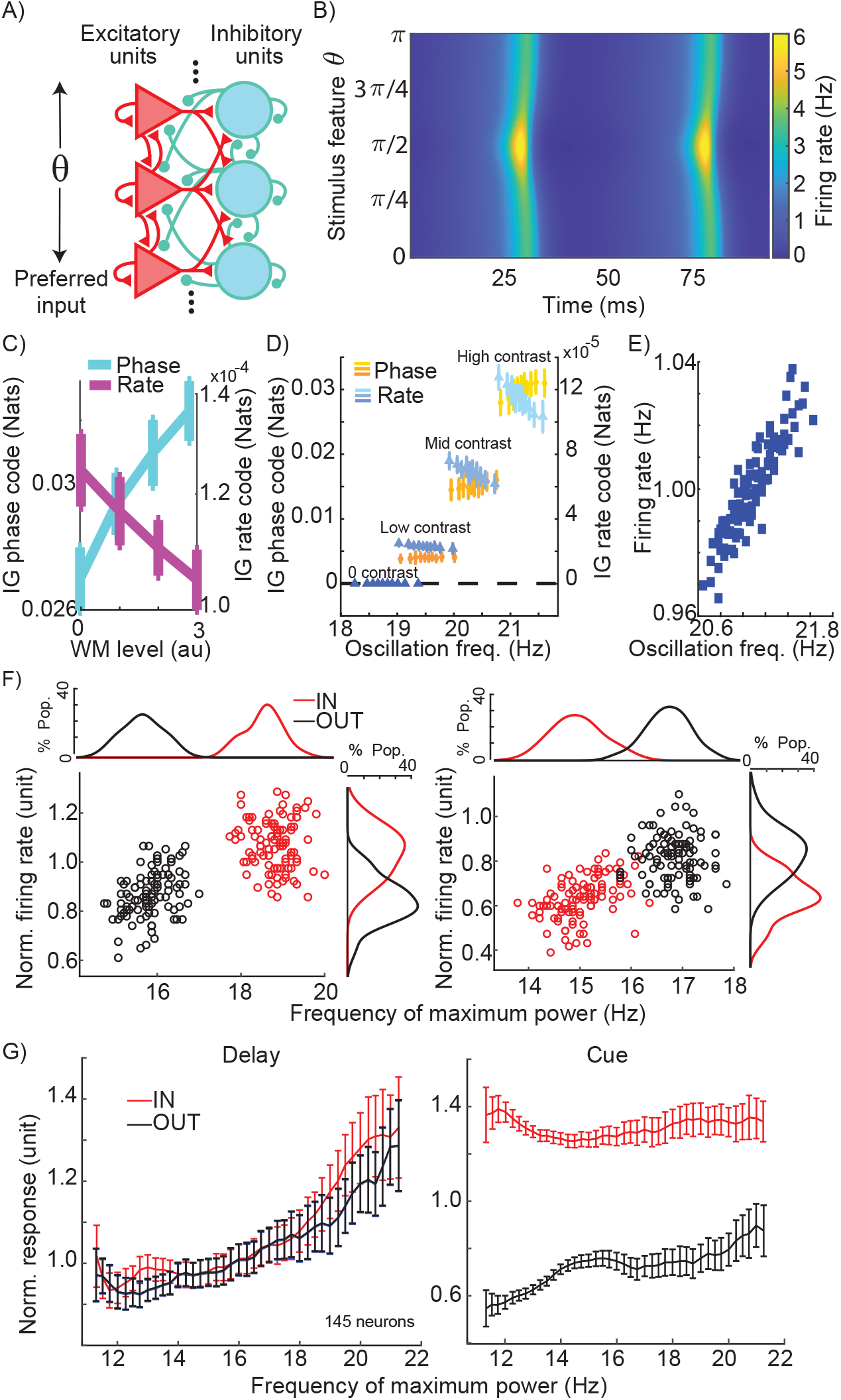
Experimental and computational dependence of firing rate, phase-coded information, and rate-coded information on changes in peak oscillation frequency. A) Schematic of dynamical neural field network architecture. Excitatory and inhibitory units are interconnected and organized to respond to different input stimuli (theta). Units also receive a global WM input (not shown); see Methods for description of connectivity weights. B) Example activity of excitatory units in the model over time, in response to an input at π/2. Excitatory units are plotted along the y-axis according to their input tuning, which ranges from 0 to π. Activity reflects both a beta-frequency oscillation across the entire population, and an earlier and stronger response of units whose preference matches the input feature (i.e., phase and rate coding). C) Information about the input stimulus feature coded by phase and rate (left and right y-axes; see Methods) in the neural field model, as a function of WM input strength. D)Phase information (shades of orange) and rate information (shades of blue) as a function oF_Contrast_ levels and oscillation frequency, for the neural field model. Data for each contrast is divided into deciles based on oscillation frequency, variability in which comes from noise in the WM input strength. Note that rate code values are several orders of magnitude smaller than phase code values (left vs. right y-axis). Error bars show standard deviation. E) Correlation between oscillation frequency and firing rate in the neural field model under conditions of noisy WM input strength. F) Relationship between frequency of peak LFP power and firing rate during the delay period for the memory IN (red) and OUT (black) conditions, for two example neurons with increased (left) or decreased (right) firing rate during the IN condition. Each dot shows the average frequency of max power and normalized firing rate for a subsample consisting of 50% of trials (n=100 subsamples per neuron). G) Average normalized response as a function of peak frequency, pooled across subsamples of trials from each of 145 V4 neurons (100 subsamples/neuron), during the IN (red) and OUT (blue) conditions, during the delay period (left) and the cue period (right). Plot shows mean±SE for all subsamples with the peak frequency indicated on the x-axis.

The model revealed that stronger WM input increased both oscillation strength and peak frequency (Fig. S12), and we hypothesized that this change in frequency could explain changes in firing rate. We tested this idea both in the model and in the experimental data. In the model, we varied the strength of the WM signal across 4 levels of stimulus input strength (resembling various levels of stimulus contrast). We then divided oscillatory cycles occurring in the respective stimulus input levels into deciles based on their oscillation frequency, and measured information encoded by phase and rate as a function of oscillation frequency (Fig. 4D). Phase information increased with increasing oscillation frequency while rate information decreased at higher oscillation frequencies, for all non-zero contrast levels (Model utility t-test for contrasts 0-3 respectively; phase code: p>0.05, >0.05, <0.01,0.001; and rate code: p>0.05, <0.05,0.01,0.0001). Importantly, we found that the network replicating the oscillatory and representational characteristics of V4 during WM shows an increased firing rate as oscillation frequency increases. Figure 4E shows how small variations in the strength of the WM signal resulted in correlated changes in oscillation frequency and firing rate (r = 0.901; linear model utility t-test significant: p<0.0001). Thus, firing rate is positively correlated with oscillation frequency, but information encoded by that rate is negatively correlated (Fig. 4C vs. 4E). To confirm that such a relationship exists in the experimental data, for each neuron we measured the peak LFP frequency and average firing rate across subsamples of trials, allowing us to test the relationship between peak β frequency and evoked firing rate within a single condition. As shown for two sample V4 neurons, such a relationship between peak frequency and average firing rate existed between the IN and OUT conditions (for a single background stimulus; Fig. 4F). For the first sample neuron, in which the peak frequency changed from 18.81 to 15.84 Hz between the IN and OUT conditions, the firing rate changed from 1.084 to 0.869, and peak frequency and firing rate were correlated across subsamples of trials (Pearson correlation, r=0.716, p<10^−32^; Fig. 4F left). Results from the second sample neuron show that this relationship remains the same even in cases where WM reduces the peak frequency (peak-frequency_IN_ = 15.041 Hz, peak-frequency_OUT_ = 16.827 Hz, FR_IN_=0.630, FR_OUT_=0.816): the correlation between firing rate and peak frequency remains positive (Pearson correlation, r=0.671, p<10^−26^; Fig. 4F right). At the population level, we looked at firing rate during the delay period across subsamples of trials for all 145 V4 neurons, sorted according to their peak frequency (Fig. 4G). As shown in figure 4G left, this relationship between peak frequency and firing rate was present across both IN and OUT conditions; more importantly, this relationship remained the same between the two memory conditions (F_Condition_ =9.649, p=0.003; F_Frequency_ =399.566, p<10^−31^; F_Interaction_ =2.935, p=0.091, ANCOVA), suggesting that the frequency of WM-induced oscillations could account for firing rate changes. Critically, a similar analysis for the visual period of the task revealed that the presence or absence of a visual stimulus (IN vs. OUT condition) creates a much larger change in firing rate, which cannot be accounted for by changes in β frequency (F_Condition_ =5.113×10^3^, p<10^−72^; F_Frequency_ =103.220, p<10^−15^; F_Interaction_ =71.427, p<10^−11^, ANCOVA) (Fig. 4G right). This relationship between frequency and firing rate was also significant within each contrast level (Fig. S14). In other words, changes in rate during the delay period may be a consequence of changes in phase locking frequency (although we cannot rule out that the firing rate is driving the frequency instead, or that changes in both rate and frequency are independently generated by a common input). We also examined the relationship between peak frequency and firing rate as a function of stimulus efficacy. Both firing rate and peak frequency vary with stimulus efficacy (Fig. S15A; two sample neurons), but there was no difference for this relationship between the IN and OUT condition (Fig. S15B). Figure S16 shows that this relationship between firing rate and peak frequency is specific to the β frequency range. While there was no overall change in average firing rate due to WM across the V4 population (see Fig. 1, Fig. S1, and related statistics), this analysis further suggests that any WM-related changes in the firing rates of individual V4 neurons could be driven by changes in oscillatory frequency.

## Discussion

Prefrontal cortex modulates sensory and motor signals in order to guide our actions based on goals and priorities maintained in WM^3,4,37,38^. Within prefrontal areas, FEF sends direct projections to extrastriate visual areas, and activity in these projections reflects the content of WM^8^. We designed a paradigm in which neurons in extrastriate area V4 are provided with bottom-up input while the top-down signal carrying WM content can be directed to the part of space represented by these neurons or elsewhere. This allowed us to examine which aspect of the sensory representation within V4 is influenced by a top-down WM signal, and to causally test the role of FEF activity in this WM-driven modulation. We found that a neural phase code representation of sensory stimuli was strongly modulated by top-down WM signals coming from the FEF, while firing rates were relatively unaffected, suggesting that representations based on the average firing rate of neurons might not be the primary way that top-down signals enhance sensory processing. Using a combination of computational modelling and experimental data analysis, we provided evidence that any changes in the average firing rate of individual neurons might be a byproduct of small changes in the frequency of the WM-induced oscillation.

Attention and WM are closely linked, although the exact nature of their relationship is not yet known ^39-44^. Behaviorally, WM can impact sensory processing, changing discrimination thresholds ^45-47^ and reducing reaction times ^48^, similar to the behavioral effects of attention. In visual areas, spatial attention and WM have several similar impacts on visual responses: shifts in RFs^8,49,50^, enhanced firing rates^8,51^, decreased variability^8,52^, and changes in inter-neuronal correlations^53,54^. Within FEF, neurons with WM-related activity show greater attentional modulation ^55^, and the same neurons in lateral PFC are involved in selection when switching between attention and WM tasks ^56^. Although behavioral training can dissociate the neural basis of WM and attention in prefrontal cortex ^57,58^, it remains likely that by default these mechanisms are linked or overlapping. We cannot rule out the possibility that the animal is allocating attention to the WM location, and that this drives the phase coding; however, the feature information we see enhanced is task-irrelevant, has no measurable impact on performance, and the monkey hasn’t been rewarded for attending to it. In the opposite direction, the task irrelevance of the stimuli may discourage attention, and this may be related to the lack of firing rate modulation. Additionally, our task involves a saccade to the remembered location. Motor planning also produces changes in visual cortex similar to those seen during attention ^59^. All three of these processes—attention, motor planning, and WM—are heavily interconnected both behaviorally and neurally ^60^, and cannot be fully dissociated in our dataset. To our knowledge, no studies have reported on whether or not phase coding of visual information is modulated by either covert spatial attention or motor preparation; further work will be needed to determine whether the phase coding we report here is specific to WM or shared with covert attention and/or motor planning.

The long^61^ and growing^62-64^ list of neural signatures of attention begs a unifying theory describing the exact mechanisms involved in generating this plethora of neural signatures. Many of these signatures are seen in both attention and WM, including enhanced visual responses^8,51^, changes in inter-neuronal correlations^53,54^, decreased variability^8,52^, and shifts in RFs^8,49,50^. In light of the present findings, we suggest that by inducing an oscillation, top-down signals allow expression of sensory representations in the form of a neural phase code: neurons emit action potentials in response to this induced oscillation with a relative timing that reflects their sensitivity. Slight changes in the frequency of the oscillation might then account for changes in the average firing rate of neurons in sensory areas, and one can imagine coherent oscillations altering the dependent and independent variability of the neurons as well. Understanding whether the β oscillations observed in our study function in the same way as the mostly gamma oscillations reported in attention studies will require a more complete understanding of the characteristics of oscillators operating in the presence and absence of visual information (see also ^53^).

A framework in which top-down signals primarily alter the phase of the spikes faces an important challenge: in communication between brain areas, LFP oscillations are not carried along with spikes down axonal projections. A phase code without its oscillatory reference frame is likely unreadable. However, studies in our lab and others have provided growing evidence that there are coherent oscillations between brain areas during WM, which could provide the shared oscillatory frame of reference required to transfer phase-coded information. For example, oscillatory coherence between FEF and inferotemporal cortex exists and predicts performance on an object WM task^65^; similarly, synchrony between PFC and V4 is also correlated with WM performance^66^. Oscillatory coherence between prefrontal and parietal areas also reflects the content of WM^67-69^. For a more complete review of findings of inter-areal coherence during WM and their relationship to performance, see ^70^. The significance of enhancing the efficacy of signals by generating a coherent signal has previously been presented in the context of communication through coherence (CTC)^71-74^, as has the idea that phase in the receiving area can influence sensitivity to incoming signals^75,76^. In addition to gating of efficacy by phase of the receiving area (as in CTC, where this gating can make a downstream area more sensitive to input from one source than another^77^), the precise timing of spikes relative to oscillations even within a coherently oscillating site (i.e. phase coding) could also be crucial when this timing is going to be gated back into signal strength using a coherent oscillation in the receiving area.

We found that WM signals allow expression of visual representations in the form of a neural phase code, indicating that prefrontal cortex could recruit sensory areas using a WM-induced oscillation. This new finding, along with the abundant evidence of coherent oscillations across brain areas during WM^70^, lead to a working hypothesis about how WM can recruit sensory areas. Consistent with sensory recruitment theories of WM^37,78-80^, these results suggest that sensory and memory signals could be preserved in sensory areas without being expressed in their average firing rate. This latent information could be expressed in the form of a phase code in response to a WM-induced oscillation, and is potentially readable by other areas that have oscillations coherent with the oscillatory frame of reference induced by WM. This proposed recruitment through coherence framework of working memory offers an explanation for how WM can recruit highly feature-sensitive sensory areas in the absence of robust firing rate changes within them.

## Materials and methods

### Experimental model details

We recorded from two male rhesus monkeys (Macaca mulatta, 12 and 16Kg). All experiments and animal procedures in this study were in accordance with the National Institutes of Health Guide for the Care and Use of Laboratory Animals and the Society for Neuroscience Guidelines and Policies. Protocols for experimental and behavioral procedures were approved by the University of Utah Institutional Animal Care and Use Committee.

### General and surgical procedures

All surgeries were performed under aseptic conditions, using standard techniques and gas anesthesia, with appropriate peri-surgical analgesia and monitoring. After the study’s conclusion, both animals remained healthy and were subsequently utilized in other research endeavors. Stereotactic surgery coordinates for the PFC and V4 chambers (20mm diameter) were performed for monkey 1, right hemisphere, at (AP 25+(2), ML 15 (±0)) and (AP -5(−1), ML 20+(2)), and for monkey 2, left hemisphere, at (AP 30+(1-2), ML 15-(1-2)) and (AP -5-(1-2), ML 20+(1-2)).

### Behavioral tasks

We programmed all behavioral tasks using the NIMH Monkeylogic toolbox (ML2) 55, on 64-bit Matlab software (The MathWorks, Inc., Natick, MA). We monitored eye position with an infrared optical eye-tracking (EyeLink 1000, SR Research, Ottawa, Canada). Visual tasks were presented on a VG248 ASUS LED monitor with a refresh rate of 144 Hz and resolution of 1920 × 1080 pixels.

#### V4 RF mapping

On a daily basis, we first identified V4 RFs using audible responses to oriented bars. Second, we presented a series of visual stimuli on a black background, to quantitatively estimate V4 RFs based on the neuron’s firing rate response. Visual stimuli were white circles (1dva diameter), 100ms on, 100ms off, pseudorandomly presented in a 7×7 grid spaced 2.5 dva between stimuli. The monkeys fixated on a central white circle (1dva diameter) throughout the trial.

#### FEF RF mapping

We estimated FEF RFs using electrical stimulation within the anterior bank of the arcuate sulcus, in biphasic microcurrent pulses (50μA) using a S88 Grass stimulator. Stimulation was performed via tungsten microelectrodes (FHC, Bowdoin, ME). FEF sites were identified based on the landing point of the evoked eye movement following stimulation with currents ≤50μA.

#### Memory guided saccade tasks

To assess the influence of WM on the representation of sensory stimuli, we used a variant of MGS task with a background stimulus (Fig. 1A). The MGS-background task is similar to classic MGS task with a task-irrelevant full field stimulus in the background. The background stimulus was an oriented grating, which could appear in one of four orientations and 4 contrasts (0% contrast is just a classic MG task). The WM cue was placed either within the overlapping RF of FEF and V4 (IN condition) or 180 degrees away (OUT condition). During FEF inactivation experiments, a classic 8 location MGS task with no background was used to assess the behavioral consequences of drug injection over space and time (Fig. 3B-E), in addition to the MGS-background task to measure information coding.

### Neurophysiological recording

We recorded the activity of 145 V4 neurons across 88 recording sessions (55 sessions Monkey E, 33 sessions Monkey O), including 66 V4 neurons during 33 FEF inactivation sessions (29 sessions Monkey E, 4 sessions Monkey O). We recorded neurophysiological activity using Neuralynx and Blackrock data acquisition systems. We digitized spike waveforms at 32 KHz, and performed offline spike sorting manually. We used single tungsten microelectrodes of 200μm diameter, with epoxylite insulation (FHC, Bowdoin, ME), and linear 16-channel arrays (Plexon, Dallas, TX). Electrodes were inserted using a hydraulic microdrive (Narishige, Japan).

#### V4 recordings

We simultaneously recorded from FEF (single electrode) and V4 (single or linear array electrodes). In this paper we only present the data from the V4 recordings.

#### FEF inactivation with V4 recording

FEF was pharmacologically inactivated through infusion of 0.5-1μL of the GABA-a agonist muscimol, using a custom microinjectrode system (described in ^81,82^). Muscimol concentration was 5mg/ml (pH 6.5 to 7). V4 activity was recorded from a site with RFs overlapping the estimated FEF RF, before and after FEF inactivation. Performance on the memory guided saccade task was used to verify FEF inactivation.

### Data analysis

#### Quantification and statistical analysis

For all analyses of V4 responses (main Figs. 1,2,4), we pooled V4 data from simultaneous FEF-V4 recording sessions with data from FEF inactivation sessions (using the V4 data prior to FEF inactivation). For the results of Figure 3, we assess the role of FEF on V4 coding using inactivation data in which we recorded from V4 before and after FEF inactivation. Evaluations of neural responses to background stimuli of varying contrast or orientation include the three non-zero contrast values. Wherever a statistical test is not specified it is Wilcoxon sign rank. P values are reported up to three decimal digits, and p values less than 0.001 are reported as p<10^−x^. The β range is defined as 14-22 Hz. The significance of the histograms was indicated using asterisks (* for p < 0.05, ** for p < 0.01, and *** for p < 0.001), while non-significant results were denoted with ‘ns’ (not significant, p > 0.05).

#### Behavioral analyses

The percentage of correct was defined as correctly completed trials divided by all completed trials (Fig. 3C). Target window size (used for % correct calculations and for rewarding the animal) was a circle of radius 2 dva for cue eccentricities <7 dva, and radius 4 dva for eccentricities >7 dva.

Reaction time (RT) was the time between cue and saccade onset. Saccade error was defined as the difference between the endpoint of the saccade and the cue location. For across-session behavioral analysis (Fig. 3D-E), RT and error were normalized within each session by dividing by the median RT/error of all before inactivation trials.

#### LFP power spectrum, spike-phase locking, and STA

The power spectral density of the local field potentials (LFPs) was calculated using the multitaper method, employing three tapers (discrete prolate spheroidal (DPSS)-Slepian sequences) for each trial and channel. For population LFP power statistics, LFP power spectrums were normalized, [(X-min)/(max-min)]. In sessions with array recordings (51/88 sessions) power calculations were performed for each channel and then averaged across all channels in that session before calculating population statistics. To quantify the reliability of spike timing relative to the LFP of the same channel, we employed the Spike-phase locking (SPL) method ^83^, which measures the consistency or locking strength of spike phases to the LFPs. This is achieved by calculating the angular summation between phases of LFPs and spike times. The amplitude of the SPL indicates the strength of spike locking to LFP phase, while the angle reflects the phases of LFPs when spikes occurred. For the spike-triggered average (STA) of the LFP, we first normalized the LFP by taking the z-score of the LFP across timepoints within 100ms of a spike for each trial, then averaged those values across trials. We concentrated on neurons with well-defined STAs. Low contrast STAs lacked a clear peak and were excluded from the analysis.

#### Rate and phase coding capacity

Our primary means of measuring coding capacity was the method developed by Panzeri and colleagues, which allowed us to quantify and compare information contained in rate and phase codes ^84,85^. This calculation of coding capacity was done in four steps. 1) First, using the FIR filter, LFPs were filtered into ten different frequency bands (1-4; 4-8; 8-12; 12-17; 17-22; 22-27; 27-35; 35-55; 65-90; 90-120). 2) Next, based on Hilbert transform, the phase of filtered LFPs were extracted. 3) Subsequently, the average of phases at the time of spike occurrence were estimated for a window of 100ms duration with 100ms shift. 4) Finally, mutual information (MI) was calculated between these average phases (phase code) or average spike rate (rate code) and different stimuli. The MI was calculated across all stimulus contrasts and orientations. For full mathematical details see ^84^. The configuration we used was: direct method, biased naive estimates and 20 bootstraps ^85^.

### Mathematical modelling methods

Details of this model were previously published in Frontiers in Computational Neuroscience ^86^. Neural field model

Our neural field model is defined by be a periodic orientation tuning domain parameter *θ*∈ [0,*π*) This neural field model is intended to represent a hypercolumn-like population with a subset of cells within the neural field preferentially responsive to a *θ*-oriented stimulus. This model has been studied in detail in a previous article by our group ^86^. The neural field model is described by *u(θ,t*), *v(θ,t*),the e- and i-activity for every *θ*-location on the ring, that solves the integro-differential equation:

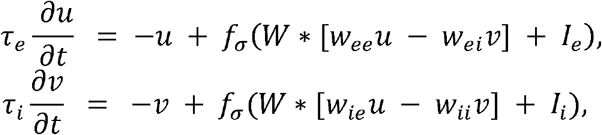

The integral convolution “*” in the above, is over the *θ*-domain, with *W*(*θ*) being the von-Mises periodic weight kernel

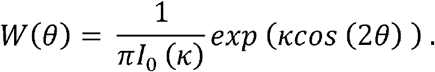

A mass-model at *θ* and *θ*’ will be connected with weight *W*(*θ*− *θ*′)*dθ*′. The parameter *κ* is the inverse-variance-like scale parameter that shapes the broadness/tightness of the distribution, and *I*_0(*k*) is the order-zero modified Bessel function of the first kind, which serves as the normalization constant. Note that the half-circle orientation tuning domain *θ* ∈ [0,*π*) necessitates a “2” factor in the weight function to be *π*-periodic. The same spatial scale *k* is used for both e- and i-cell populations.

The *f*_*σ*_ (1) function defines the output firing rate of each population as a function of its input *I*---an F-I curve. We have used a sigmoidal-shaped F-I curve defined as the inverse mean first passage time, plus a 5ms refractory period, of a leaky integrate an fire LIF model neuron driven by uncorrelated Gaussian white noise *σ ξ* (*t*) with standard deviation *σ* (see for example ^86,87^:

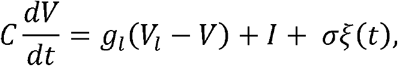

with capacitance *C*= 1 micro-Farads, spike threshold voltage *V*_***t***_ = -50mV, and reset and leak voltages *V*_***r***_ = *V*_***l***_ = −65mV (all parameters are listed in Table S1). With these parameters, *I*= 1nA of current induces the membrane voltage to approach spike threshold in the absence of noise. Increasing noise parameter *σ* has the effect of reducing the overall gain of the F-I curve (see ^86^, for more details on this model).

Three types of external inputs were given to the neural field: working memory inputs, stimulus inputs, and random inputs. WM inputs are uniform current inputs, added to *I*_***e***_ and *I*_*i*_, over the entire neural field (equal for all *θ*-values). These uniform inputs raised very slightly the mean firing rate and oscillation frequency and represent an WM-like or attentional-like enhancement of hypercolumn activity. Stimulus inputs are orientation-tuned given by Von Mises-like distribution functions

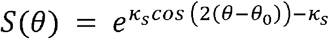

in which the peak strength (set to unity) of the stimulus located at orientation *θ*_**0**_. We fix *θ*_**0**_ = *π*/2--a 90-degree (vertical) orientated stimulus, without loss of generality. Finally, to capture the temporal variations in network oscillations observed in real cortical tissues, on simulations in which we assessed sensory coding (see below), we included slow-timescale Ornstein-Ulenbeck noise *y*(*t*) to both e- and i-cell input currents *I*_***e***_ and *I*_*i*_ globally to the entire network (uniformly across all *θ*-values). The dynamics of *y* are given by the stochastic differential equation

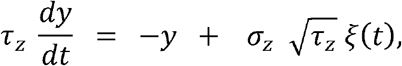

where *ξ*(*t*) is uncorrelated zero-mean unit-variance gaussian white noise. This equation results in a normal stationary distribution of *y*-values, with zero-mean, and standard deviation *σ*_***z***_, and a temporal autocorrelation decay timescale *τ*_***z***_ = 50ms, so that the network oscillations, which were typically in the 20 Hz range (50 ms oscillation cycles), showed robust cycle to cycle variability but little long-timescale multi-cycle correlation.

In the absence of any external input, we set neural field model to be very near the a supercritical Hopf instability (see ^86^) in which additional current above a current threshold *I*^***\****^, elicited oscillations with amplitude emerging continuously from zero, and oscillation frequency in the *β*-band around 18-20 Hz. From this *I*^***\****^ parameter starting point we ran simulations from over four levels of WM input (uniform current) and four levels oF_Orientation_-selective stimulus input (contrast levels), starting from zero. We call these WM 0,1,2,3 levels, and contrast levels 0,1,2,3. Altogether, the input to cells can be represented by

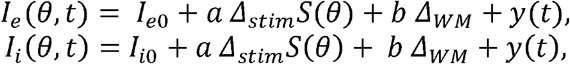

where *Δ*_***stim***_ and *Δ*_***WM***_ are the current increments for the respective input levels of stimulus *a* = 0,1,2,3 and WM *b* = 0,1,2,3. In addition to the input current changes that occur for our model, it is common to accept that increased stimulus input comes with increased input current fluctuations. We modeled this by adjusting the *σ* -parameter the F-I curve as a function oF_Contrast_ input:

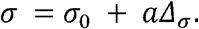

#### Coding performance

The phase- and rate-based responses of the neural field model can be used to discriminate the input stimuli. We have chosen to discriminate the neural field model responses at the neural field locations *θ* = *π*/2 and *π*/4. We computed the coding performance for two competing codes: a rate code (i.e., a spike count code) and a phase code. To define the phase variable in the phase code, we examined the proxy LFP signal formed by averaging e-cell rate responses over the entire field domain. We derived a phase angle *φ* (*t*) ∈ [− *π, π*] of oscillation via the Hilbert transform of this LFP signal. After segmenting the simulation run time into oscillation cycles *φ* (*t*) ∈ [− *π, π*], for *t*∈ [0,*T*], where *T* = 1/*f* is oscillation period. The mean rate response is simply 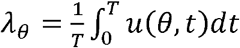, from which we assume Poisson-distributed *n* number of spikes are emitted:

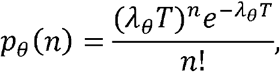

which constituted the rate code distribution.

The phase code distribution is obtained from the rate response *u*(*θ,t*), by using a change-of-variables between time and phase *t*= *g*(*φ*), where *g*(*φ*) is the inverse of the Hilbert phase angle:

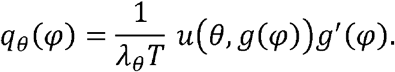

Using the spike count and phase distributions as the basis of the rate and phase codes, respectively, we computed two different measures of coding performance. First, to measure the amount of information gained from the code at *θ* = *π*/2, given one assumes the data are distributed according to *θ* = *π*/4, we computed the information gain rate (IG)--the Kullback Libler divergence (in natural units of information, nats):

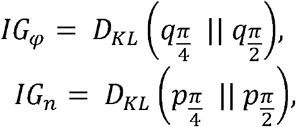

where 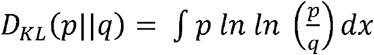. Second, we computed the mutual information (MI) between stimulus feature *φ* = *π*/2 or *π*/4, and the phase data *φ*. We assumed the two stimuli were equally likely on a given “trial” in which case the probability of each stimulus was 1/2. The phase mutual information *MI*_*φ*_ is then defined as the Kullback Libler divergence *D*_*KL*_ (using log-base-two, in this case) between the pooled distributions 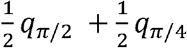 to the product distribution *q*_*π*/2_ *q* _*π*/4_ ;and similarly, for the rate codes:

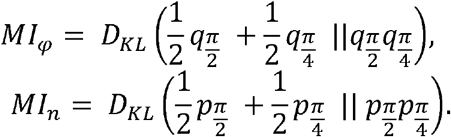

## Data availability

The code for mathematical modelling are publicly available at https://osf.io/dhcr2/. The full dataset is publicly available at https://gin.g-node.org/KClark/Parto_eLife_2024 (doi: 10.12751/g-node.6icjl8). Further information for data and resources should be directed to Lead Contact, Behrad Noudoost.

## Acknowledgements

This work was supported by NIH grants EY026924 & NS113073 to B.N, and an Unrestricted Grant from Research to Prevent Blindness, New York, NY, to the Department of Ophthalmology & Visual Sciences, University of Utah.

## Author contributions

Conceptualization, M.P., I.V., and B.N.; methodology, I.V., K.C., and B.N.; software, M.P., M.Z.; formal analysis, M.P. and M.Z., B.N.; modeling: W.N.; writing – original draft, M.P. and M.Z., K.C., B.N.; writing – review & editing, all authors; visualization, M.P., W.N., K.C.; supervision, B.N.; funding acquisition, B.N.

## Supplemental information

Document S1. Figures S1-S16 and Table S1.

