## Supplementary material for "Prefrontal working memory signal controls phase-coded information within extrastriate cortex": Document S1. Figures S1-S16 and Table S1.

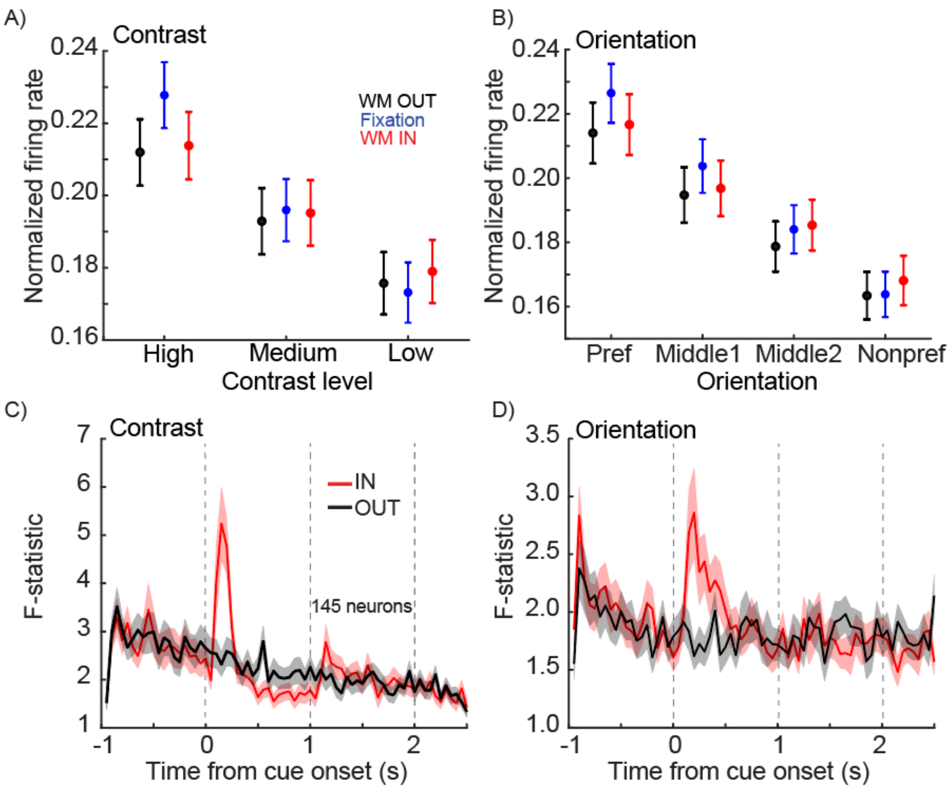


**Figure S1) V4 neurons' sensitivity to stimulus contrast and orientation is not altered by WM.** To determine whether the top-down WM signal interacted with the bottom-up sensory input to modulate V4 firing rates, we compared the firing rate of V4 neurons across different background stimuli and WM conditions. A, B) Mean normalized firing rate of 145 V4 neurons during fixation (blue), the delay period of working memory IN (red), and the delay period of working memory OUT (black) for different background stimulus contrasts (A) and orientations (B). Orientations were sorted by the neuron's preference, as measured by firing rate during the fixation period (Pref, highest firing rate; Nonpref, lowest firing rate). Error bars show standard error of the mean (SEM). C, D) Time course of averaged F-statistic values based on one-way ANOVA for discrimination between three different contrasts (C) or between four different orientations (D) for IN (red) and OUT (black). Sensory discrimination peaked just following cue onset during the IN condition. However, there was no significant difference in discriminability for different background stimuli between the IN and OUT conditions during the delay period, either for stimulus contrast (Contrast: $\text{discriminability}_{IN}$= 1.995±1.997, $\text{discriminability}_{OUT}$= 1.910±1.866, p=0.139) or for orientation (Orientation: $\text{discriminability}_{IN}$= 1.771±1.613, $\text{discriminability}_{OUT}$= 1.814±1.508, p=0.269). Thus, the top-down WM signal did not alter V4 neuron's firing rates in response to different bottom-up visual input.

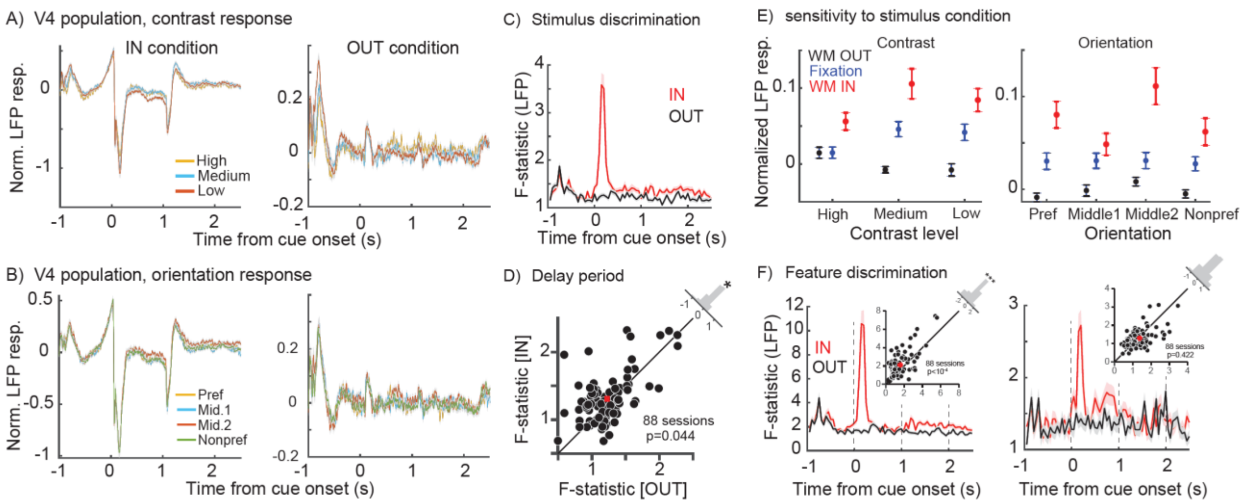


**Figure S2) V4 LFP responses to visual stimuli are altered by WM.** A, B) Mean normalized LFP response of the population of neurons over time for three different contrasts (low, medium, and high contrast, A) and four different orientations (preferred, nonpreferred, and middle 1 & 2 orientations, B) for the IN condition (left) and OUT (right) conditions. C) Time course of mean F-statistic values for the LFP across 88 sessions, based on a one-way ANOVA for discrimination between 12 stimulus conditions for the IN (red) and OUT (black) conditions. D) Scatter plot of F-statistic for an ANOVA across all 12 background stimuli, averaged in the last 700ms of the delay period for each session, for the IN (red) vs. OUT (black) conditions. Histogram in the upper right shows the distribution of change in F-statistic (OUT-IN) across sessions. The ability of the V4 LFP to discriminate between all 12 background stimuli was greater for the IN condition ($discriminability_{IN}$=1.309±0.359, $discriminability_{OUT}$=1.224±0.338, p=0.044). E) Mean normalized LFP response of the population during fixation (blue), the delay period of working memory IN (red), and the delay period of working memory OUT (black) for different background stimulus contrasts (left) and orientations (right). Orientations were sorted by the neuron's preference (as measured by firing rate of neurons recorded simultaneously with the LFP) during the fixation period (Pref, highest firing rate; Nonpref, lowest firing rate). Error bars show standard error of the mean (SEM). F) Time course of averaged F-statistic values based on one-way ANOVA of LFP responses for discrimination between three different contrasts (left) or between four different orientations (right) for IN (red) and OUT (black). There is a significant difference in LFP discriminability for different background stimuli between the IN and OUT conditions during the delay period, for stimulus contrast (left, Contrast: $discriminability_{IN}$= 2.095±1.395, $discriminability_{OUT}$= 1.591±1.041, $p<10^{-4}$) but not for orientation (right, Orientation: $discriminability_{IN}$= 1.298±0.505, $discriminability_{OUT}$= 1.368±0.604, p=0.422). Red dots on scatter plots show population mean.

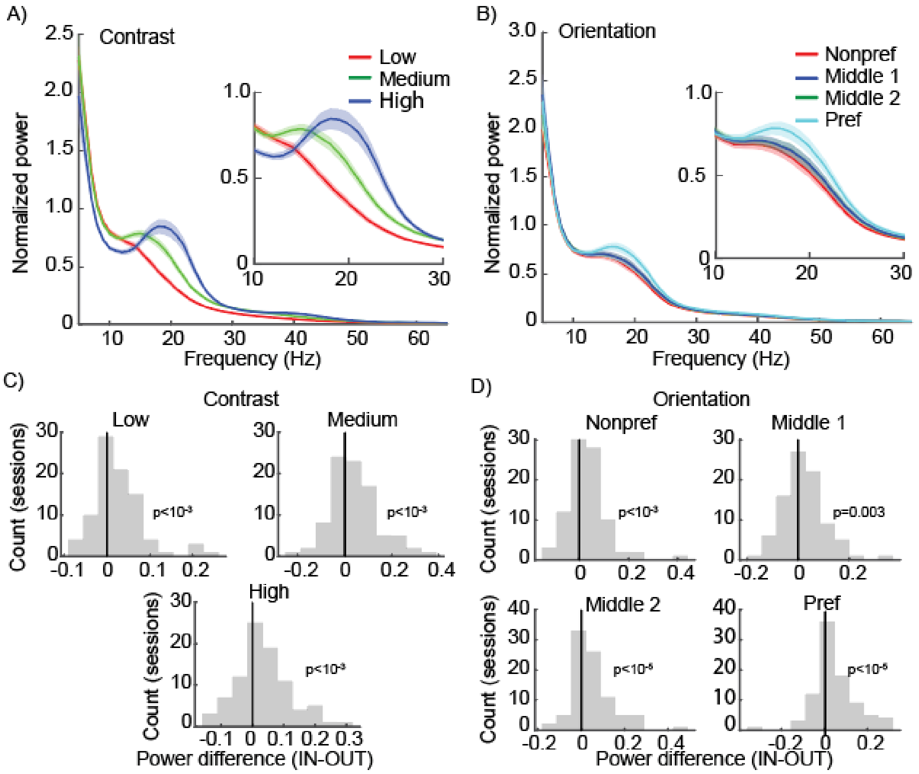


**Figure S3) LFP αβ power and frequency reflect stimulus properties and content of WM.** We examined how the top-down WM signal and bottom-up visual input altered the LFP power spectrum. A, B) Average LFP power spectrum during the delay (n = 88 sessions), for the memory IN condition, in response to backgrounds with three different contrasts (A) and different orientations (B, sorted according to the preferred orientation of neurons recorded at the same time as the LFP). More effective background stimuli (higher contrast, preferred orientation) stimuli were associated with a larger peak in the β range, with the frequency of peak β LFP power increasing from ~14 Hz for low contrast to ~19 Hz for high contrast stimuli. Inset shows 10-30Hz range. Shaded areas show standard error of mean (SEM). The β range LFP thus reflected the properties of the visual stimulus. There is a statistically significant change in peak frequency with contrasts (n = 88 sessions; one-way ANOVA, $F_{Contrast}=10.72$, $p<10^{-4}$) and with orientations (one-way ANOVA, $F_{Orientation}=3$, $p=0.030$). C, D) Histograms show the distribution of LFP power differences (IN – OUT) in the αβ range (10-20 Hz) across sessions for each contrast (C, Contrast: $\Delta Power_{low}=0.026\pm0.060$, $\Delta Power_{middle}=0.041\pm0.102$, $\Delta Power_{high}=0.036\pm0.086$) or orientation (D, Orientation: $\Delta Power_{nonpref}=0.032\pm0.080$, $\Delta Power_{middle1}=0.022\pm0.084$, $\Delta Power_{middle2}=0.049\pm0.094$, $\Delta Power_{pref}=0.048\pm0.104$). αβ LFP power significantly increased in the IN condition for all background stimuli (p-values shown on plots; Wilcoxon signed-rank). Thus, a WM signal directed toward the RF of the V4 LFP site significantly increased β power across all contrasts and orientations of background stimuli.

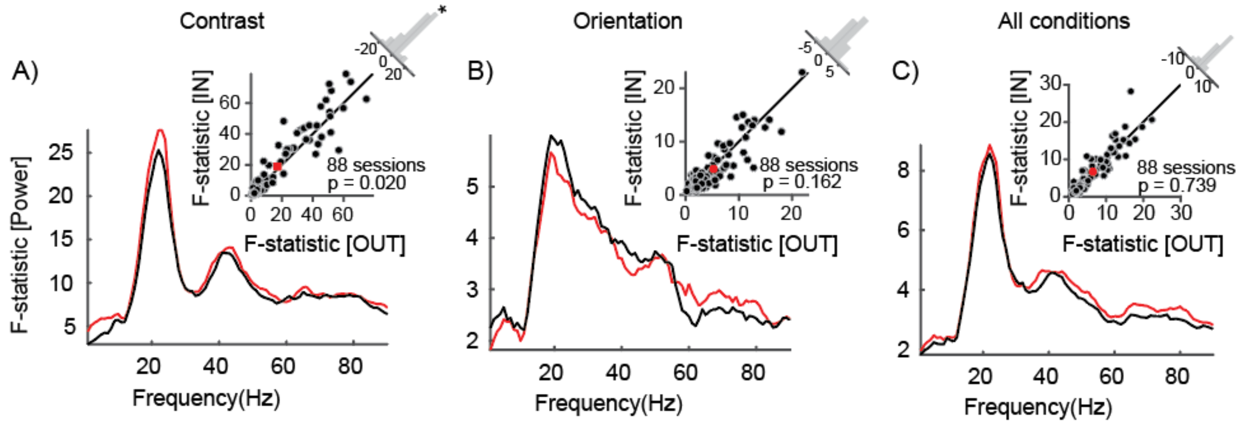


**Figure S4) V4 LFP β power sensitivity to stimulus contrast is altered by WM.** A-C) Mean F-statistic values of the power of delay period, across frequencies, based on a one-way ANOVA for discrimination between different contrasts (A), orientations (B), or 12 stimulus conditions (C) for the IN (red) and OUT (black) conditions. Scatter plots show the F-statistic averaged in β band for each session, for the IN vs. OUT conditions. Histogram in the upper right shows the distribution of change in β F-statistic (OUT-IN) across sessions. The V4 β power discriminates between contrasts for the IN vs. OUT conditions ($discriminability_{IN}$=18.894±21.344, $discriminability_{OUT}$=17.157±19.620, p=0.020; A) but there was no significant V4 β power discrimination between orientations ($discriminability_{IN}$=4.876±4.356, $discriminability_{OUT}$=5.244±4.298, p=0.162; B), or between all 12 stimuli of various contrasts and orientations ($discriminability_{IN}$=6.654±5.737, $discriminability_{OUT}$=6.428±5.170, p=0.739; C). Red dots on scatter plots show population mean.

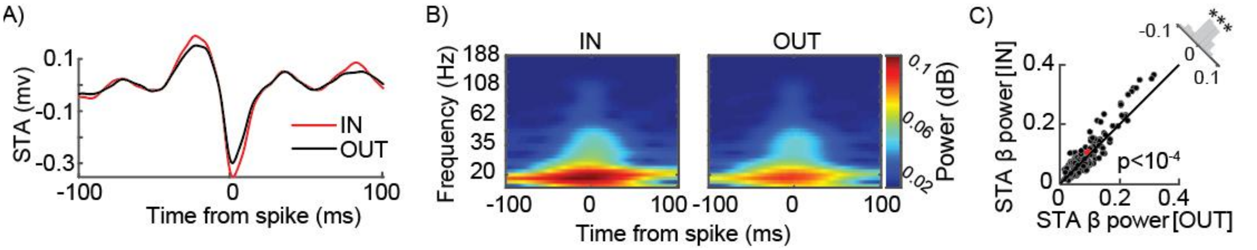


**Figure S5) β power of the LFP STA is modulated by working memory.** We calculated the spike-triggered average (STA) of the LFP during the delay period, and compared its spectral properties between memory conditions. A) STA of the normalized LFP during the delay period, for the population of 145 V4 neurons, for memory IN (red) and OUT (black) conditions. Notice that the peak and trough values are greater in the IN condition. B) Power of the LFP STA across frequency and time from the spike for memory IN (left) and OUT (right) conditions. The STA power is primarily in the lower frequencies. C) Scatter plot of STA power, averaged in β range, for IN versus OUT conditions across 88 sessions. Histogram shows the distribution of the difference in STA β power across neurons (OUT-IN). There was significantly greater β power in the STA for the IN condition (STA β $power_{IN}$=0.104±0.076, STA β $power_{OUT}$=0. 092±0.063, $p<10^{-4}$, Wilcoxon signed-rank). Red dot on scatter plot shows population mean.

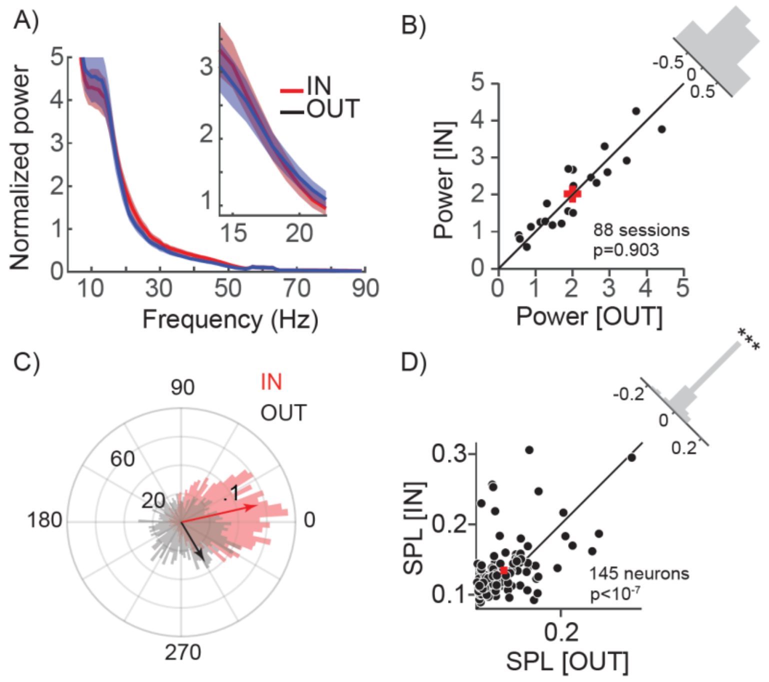


**Figure S6) WM increases V4 β oscillatory power and SPL in the MGS task with no background.** Analyses in this figure are for the 0% contrast condition, equivalent to a standard no-background MGS task. A) Mean power spectrum of population of the V4 LFPs (88 sessions) during the delay period for the IN (red) vs. OUT (black) conditions in 0% contrast background condition. Inset shows the power spectrum in β range. B) Scatter plot of power spectrum averaged in the β range for each session, for the IN vs. OUT conditions. β power shows no significant difference between IN and OUT conditions (n = 88 sessions; $Power_{IN}$=1.957±0.0.651, $Power_{OUT}$=1.818±0.733, p=0.006, Wilcoxon signed-rank). Histogram in the upper right shows the distribution of change in power (OUT-IN) across sessions (**, p <.01). C) The distribution of spikes generated by a sample V4 neuron across various phases of β oscillations during the delay period. Arrows show the average of phase distributions for the IN (red) and OUT (black) conditions; length of arrow corresponds to SPL. For this sample neuron, SPL changed from 0.088 for the OUT to 0.142 for the IN condition. D) Scatter plot of SPL in the β range for each neuron, for the IN vs. OUT conditions. SPL was significantly higher in the IN condition (n=145 neurons, $SPL_{IN}$=0.135±0.048, $SPL_{OUT}$=0.119±0.055, $p<10^{-7}$, Wilcoxon signed-rank). Histogram in the upper right shows the distribution of change in SPL (OUT-IN) across neurons (***, p <0.001). Red dots on scatter plots show population mean.

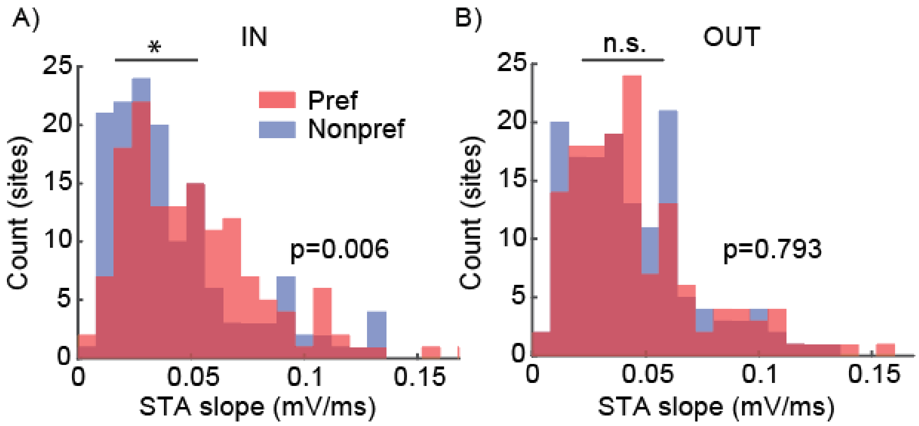


**Figure S7) Spike-LFP relationship depends on stimulus orientation and WM condition.** We tested how the slope of the LFP STA around the time of spikes varied based on background stimulus orientation and memory condition. A, B) Histogram of the distribution of slopes (abs $(V_{peak} - V_{trough}) / (time_{peak} - time_{trough})$) of the STA of the normalized LFP across neurons, for the preferred orientation (Pref, red) and nonpreferred orientation (Nonpref, blue) of the background stimulus (calculated based on neuron's firing rates during fixation), for the IN (A) and OUT (B) conditions. For the IN condition, the slope of the LFP STA was significantly different between the preferred and nonpreferred background stimuli (IN condition: $Slope_{Pref}$=0.051±0.003, $Slope_{Nonpref}$=0.043±0.003, p=0.006, Wilcoxon signed-rank). For the OUT condition, there was no difference in the slope of the LFP STA based on the background stimulus (OUT condition: $Slope_{Pref}$=0.046±0.003, $Slope_{Nonpref}$=0.044±0.004, p=0.793, Wilcoxon signed-rank). This follows the pattern of results reported for changes in the STA slope for high vs. low contrast background stimuli in Figure 2D.

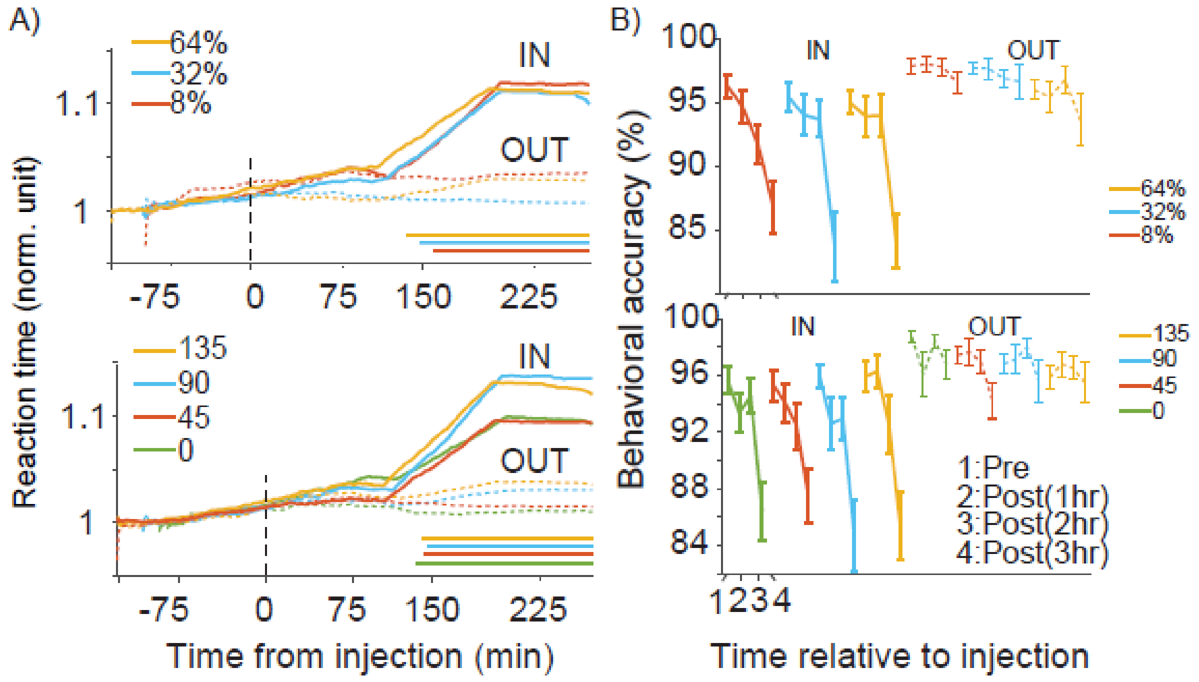


**Figure S8) FEF inactivation disrupts behavior irrespective of background stimulus and does not alter rate-based discriminability between stimuli.** A) Normalized reaction times for the memory IN (solid line) and OUT (dashed line) for different contrast (top) or orientation (bottom) conditions, over time relative to the FEF inactivation. Horizontal bars indicate times with a significant difference between IN and OUT. B) Average behavioral performance across sessions, at different stimulus background over time following FEF inactivation (pre-inactivation, 1, 2, and 3 hours after inactivation).

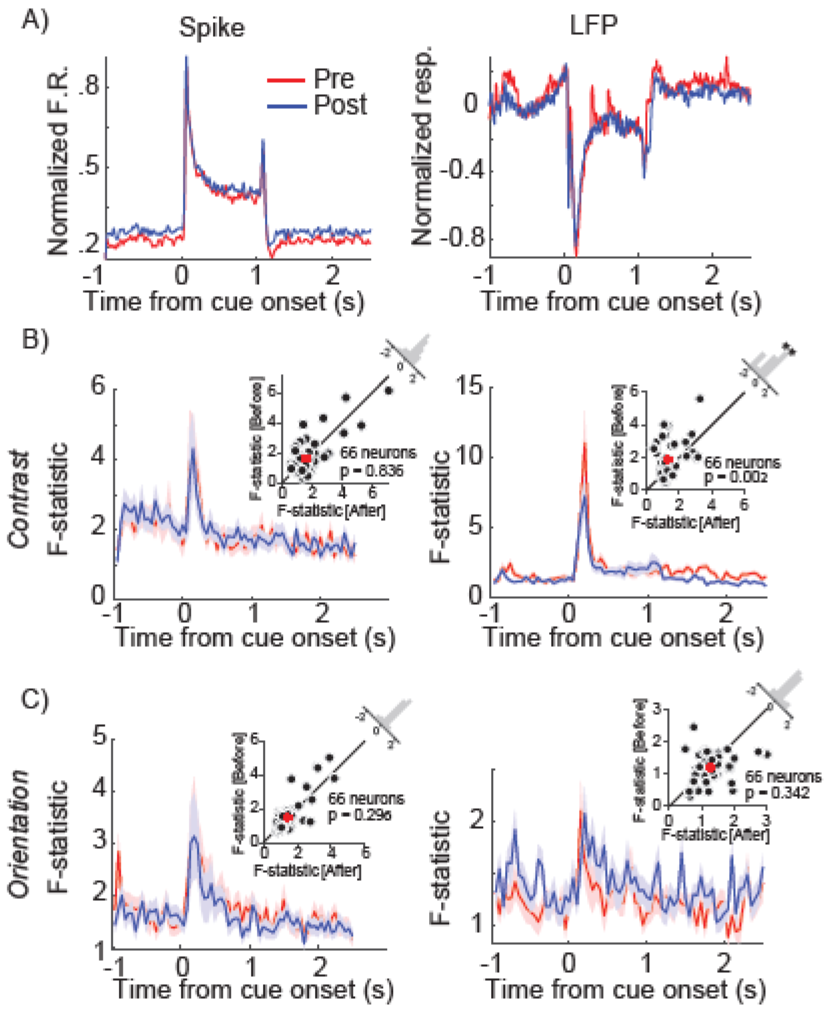


**Figure S9) FEF inactivation disrupts behavior irrespective of background stimulus and does not alter rate-based discriminability between stimuli.** A) Mean firing rate (left) and mean LFP response (right) of the population of neurons of the IN condition over time, before (Pre, red) and after (Post, blue) FEF inactivation. B, C) Time course of mean F-statistic values across 66 neurons, for firing rate (left) and mean LFP response (right), based on a one-way ANOVA for discrimination between different contrasts (B) or orientations (C) for before (Pre, red) and after (Post, blue) FEF inactivation. Insets show scatter plot of F-statistic averaged in the last 700ms of the delay period for each neuron, for the Pre vs. Post conditions. Histogram in the upper right shows the distribution of change in F-statistic (Post-Pre). Red dots on scatter plots show population mean. FEF inactivation did not alter contrast or orientation discrimination based on firing rates, or orientation discrimination based on the LFP, but did reduce contrast discrimination based on the LFP.

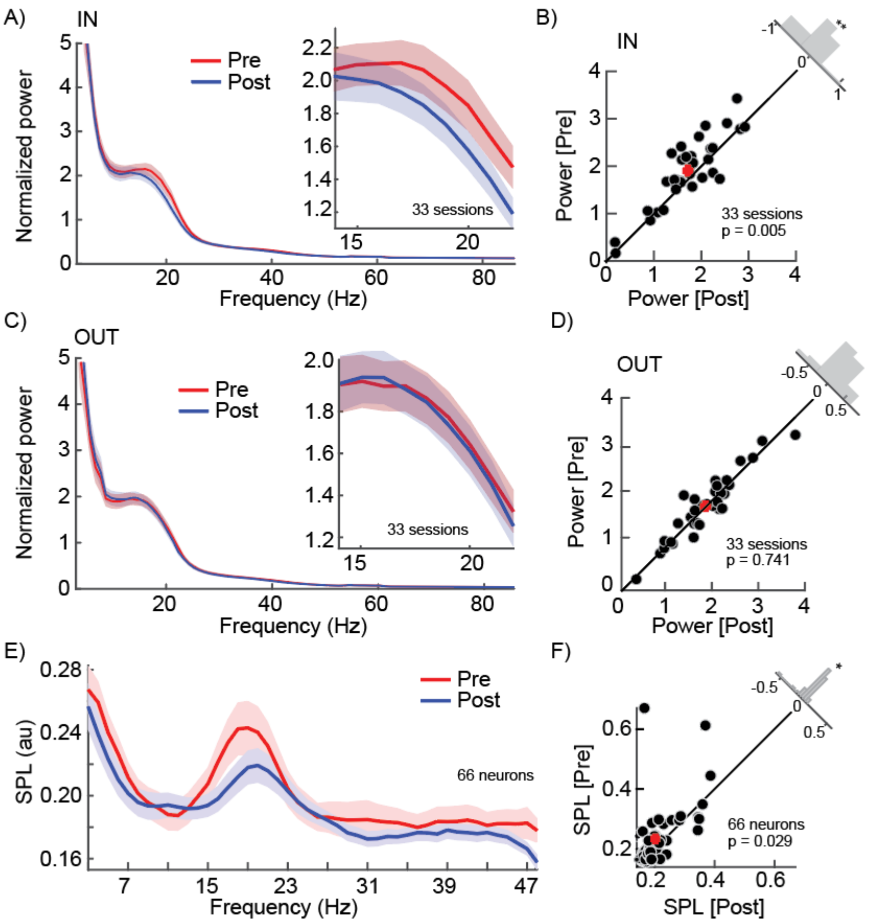


**Figure S10) FEF inactivation reduces WM-driven β LFP power and SPL in V4.** To test whether FEF activity was responsible for the WM-driven boost in V4 β LFP power and SPL, we compared V4 LFP power and the synchronization of spikes to the LFP oscillation at the same sites before vs. after pharmacological FEF inactivation. A) Normalized V4 LFP power spectrum during the delay period of the IN condition, before (Pre, red) and after (Post, blue) FEF inactivation. Inset shows power in β range (14-22 Hz). Shaded areas show standard error of mean (SEM). B) Scatter plot of V4 β LFP power for the IN condition, Pre versus Post FEF inactivation across 33 sessions. Histogram in upper right shows the distribution of the β power difference across sessions. There was significantly more β power for Pre inactivation condition (β power$_{Pre}$=1.907±0.228, β power$_{Post}$=1.720±0.296, p=0.005, Wilcoxon signed-rank). C, D) Same as A, B but for OUT condition. There was no change in V4 β LFP power before vs. after FEF inactivation for the OUT condition (β power$_{Pre}$=1.767±0.242, β power$_{Post}$=1.751±0.271, p=0.741, Wilcoxon signed-rank). E) Average SPL between V4 spikes and the phase of LFP oscillations as a function of LFP frequency during the delay period, for Pre (red) and Post (blue) FEF inactivation during the IN condition. Shaded areas show standard error of mean (SEM). F) Scatter plot of SPL in the β range for each neuron, Pre versus Post FEF inactivation. Histogram in the upper right shows the distribution of change in SPL (Post-Pre) across neurons. V4 β SPL was significantly lower following FEF inactivation (SPL$_{pre}$=0.227±0.093, SPL$_{post}$=0.202±0.054, p=0.005, Wilcoxon signed-rank). Red dots on scatter plots show population mean. These results demonstrate that FEF activity is necessary for the WM-driven boost in V4 β LFP power and SPL observed when remembering a location in the shared RF.

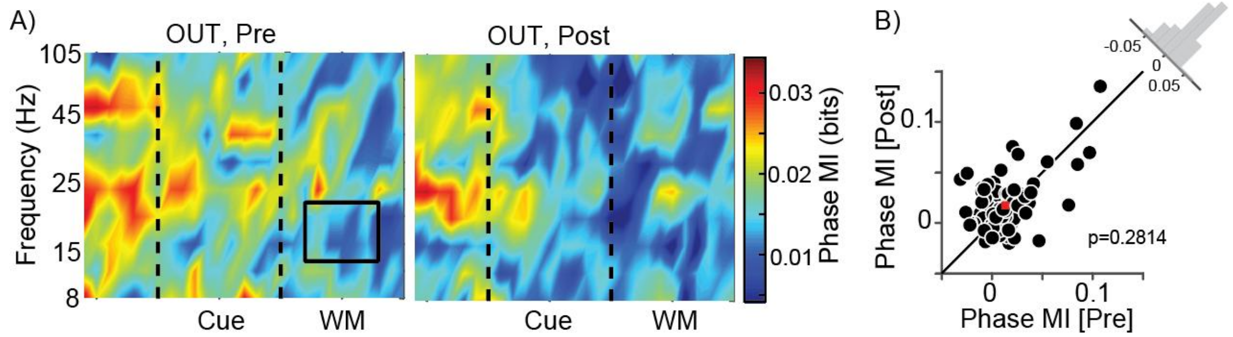


**Figure S11) FEF inactivation does not alter phase coding in the OUT condition.** A) Heatmap shows phase coding (MI, colorbar) over time and frequency for 66 V4 neurons, for the OUT condition, before (left) and after (right) FEF inactivation. Black rectangle indicates time and frequency range considered in (B): 14-22Hz, 200-800ms after start of delay period. B) Scatter plot of β phase MI during the delay period (Black rectangle in A) of the OUT condition for each V4 neuron, before vs. after FEF inactivation. Red square shows population mean. The histogram in the upper right shows the distribution of differences in MI (Pre-Post) across neurons; red dot shows population mean. There was no significant change in β phase MI of the OUT condition following FEF inactivation (n = 66 neurons; Pre, phase MI = 0.017±0.003 bits and Post, MI = 0.013±0.003 bits ; p = 0.281, Wilcoxon signed-rank).

**Table S1:** Parameter values utilized in the neural mass and neural field models. See Methods section for definitions of each parameter. For details on parameter selection and exploration of parameter space, see (Nesse et al., 2023).

| **Parameter** | **Value** |
|---|---|
| $\mathrm{V_t}$ | -50 mV |
| $\mathrm{V_r}$ | -65 mV |
| $\mathrm{V_l}$ | -65 mV |
| $\mathrm{C}$ | 1 mF |
| $\tau_i$ | 16 ms |
| $\tau_e$ | 5 ms |
| $g$ | 1/15 mS |
| $\sigma_0$ | 3 nA |
| $I_{e0}$ | -1.71 nA |
| $I_{i0}$ | -1.21 nA |
| $\Delta_{stim}$ | 0.018 nA |
| $\Delta_{WM}$ | 0.015 nA |
| $\Delta_{\sigma}$ | 0.03 nA |
| $w_{ee}$ | 0.9 |
| $w_{ii}$ | 1.9 |
| $w_{ie}$ | 1 |
| $w_{ei}$ | 2 |
| $\kappa$ | 5.0625 |
| $\kappa_s$ | 20 |
| $\sigma_z$ | *0.02* |
| $\tau_z$ | *50 ms* |

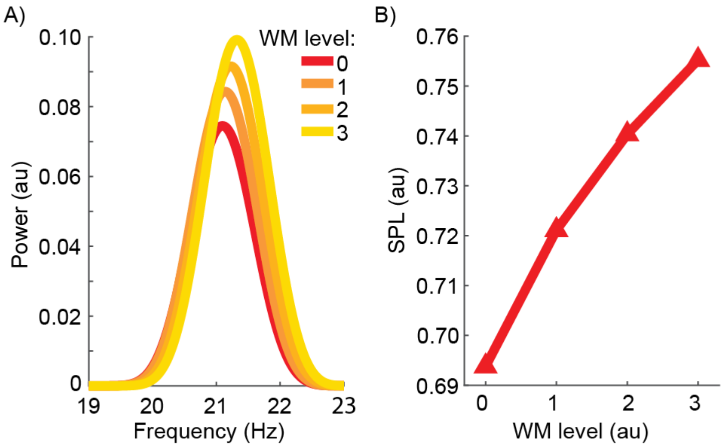


**Figure S12) Neural field model replicates key experimental findings of WM's impact on LFP power and spike timing.** We built a neural field model of visual areas receiving bottom-up sensory and top-down WM input signals (Nesse et al., 2023). This model exhibits β oscillations and locking of spikes to these oscillations; here we examine how oscillatory power and SPL change with increasing WM input strength, and compare to previously reported experimental results (where memory IN vs. OUT correspond to different WM input strengths). A) Model LFP power across frequencies, as a function of WM input strength (color). The peak LFP power and frequency of maximum power both increase with greater WM input. Increases in β LFP power with WM input have been experimentally observed in MT (Bahmani et al., 2018), and in V4 (Fig. 1I&J). B) Model SPL in the β band as a function of WM input strength. β SPL increases with greater WM input, as experimentally observed in V4 (Fig. 2B). Thus, the model replicated key findings regarding the effect of WM on LFP oscillatory power and SPL in visual areas.

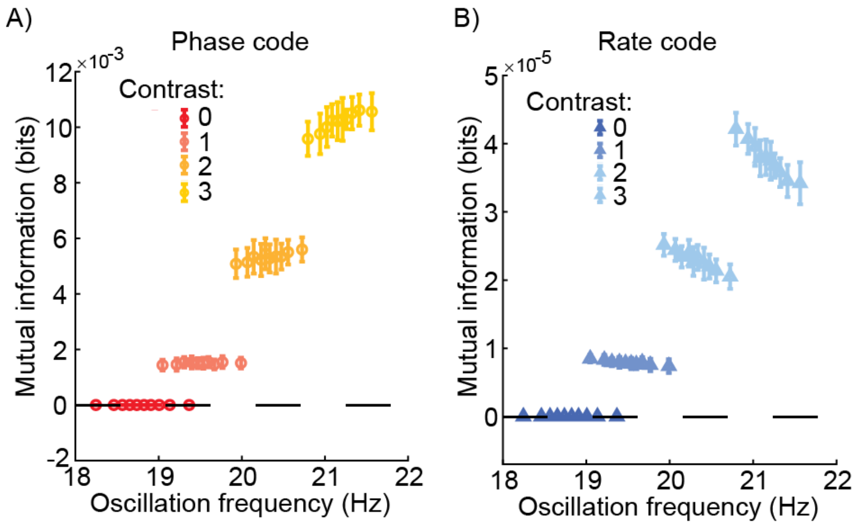


**Figure S13) Model-derived mutual information for phase and rate codes.** To test the relationship between information content and oscillation frequency in our model, we introduced noise into the strength of our WM input, and separated data for each oscillatory cycle with a given stimulus input strength based on oscillation frequency. We then calculated the mutual information between the two stimuli over four contrast levels, segmented by oscillation frequency deciles, using the phase code (A) or rate code (B). The results calculated using mutual information replicate the pattern of results reported using the information gain measure in figure 4E: phase coded information increased with increasing oscillation frequency while rate coded information decreased at higher oscillation frequencies.

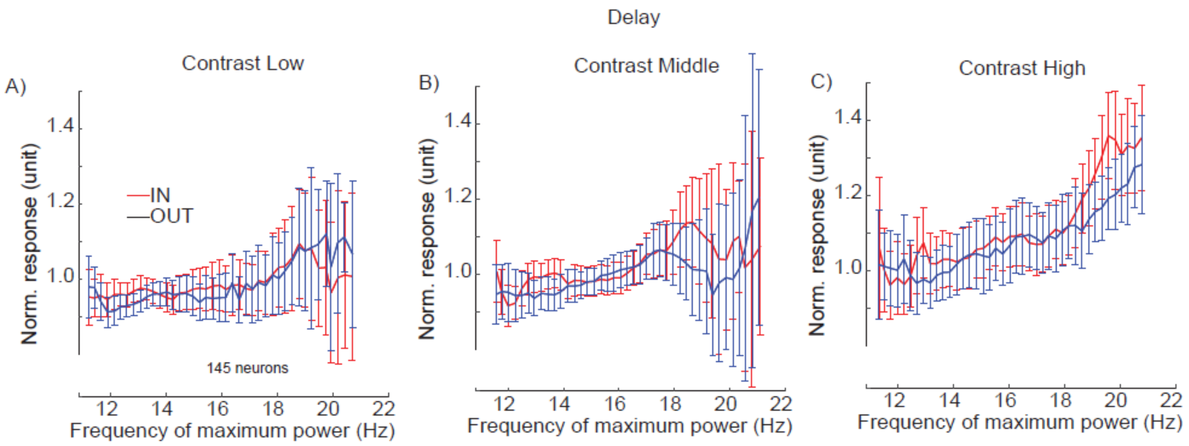


**Figure S14. Dependence of firing rate on changes in peak oscillation frequency for each contrast.** Average normalized response as a function of peak frequency, pooled across subsamples of trials from each of 145 V4 neurons (100 subsamples/neuron), during the IN (red) and OUT (blue) conditions, during the delay period for each contrast (A-C). Similar to the delay period plot in figure 4G, but separately for each contrast level. Plot shows mean±SE for all subsamples with the peak frequency indicated on the x-axis. The relationship between peak frequency and firing rate was present across both IN and OUT conditions and remained the same between the two memory conditions for each individual contrast (low contrast $F_{Condition}$=0.03, p=0.867; $F_{Frequency}$=141.86, p<$10^{-18}$; $F_{Interaction}$=10.70, p=0.002, ANCOVA; middle contrast $F_{Condition}$=7.18, p=0.009; $F_{Frequency}$=96.76, p<$10^{-14}$; $F_{Interaction}$=0.13, p=0.716, ANCOVA; high contrast $F_{Condition}$=12.51, p=0.001; $F_{Frequency}$=333.74, p<$10^{-29}$; $F_{Interaction}$=7.91, p=0.006, ANCOVA).

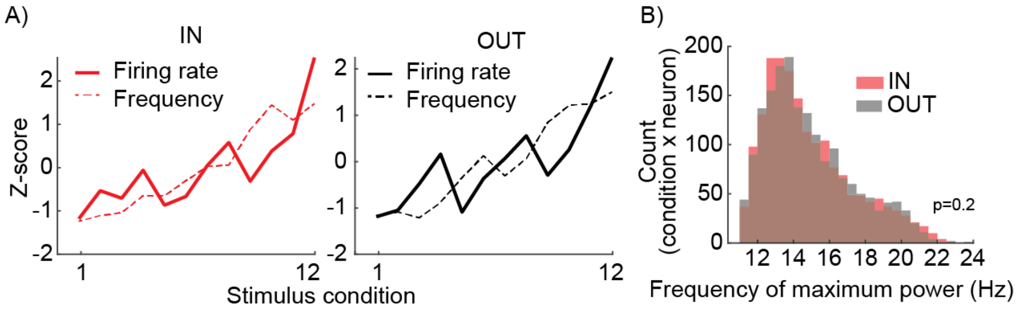


**Figure S15) Firing rate and frequency of maximum power as a function of stimulus efficacy.** In our experimental data, we looked for whether the observed relationship between oscillation frequency and firing rate varied for different combinations of background stimuli and memory condition. A) Changes in firing rate (solid line) and frequency of the maximum β LFP power (dashed line) as a function of background stimulus for memory IN (left) and OUT (right) conditions, during the delay period. Z-score reflects the difference from the average firing rate or average frequency of maximum power across all 12 stimulus conditions. The 12 background stimuli (orientation and contrast combinations) were sorted according to the firing rate of the neuron during the fixation period (weakest stimulus #1 to strongest stimulus #12). B) Histogram shows the distribution of peak LFP frequencies during the delay period, calculated separately for the 12 stimulus conditions for each neuron, for the memory IN (red) and OUT (grey) conditions ($PeakFreq_{IN}$=14.853±2.461, $PeakFreq_{OUT}$=14.885±2.427, p=0.210, Wilcoxon signed-rank). These results show that the relationship between LFP frequency and firing rate in V4 is consistent across different bottom-up inputs and memory conditions.

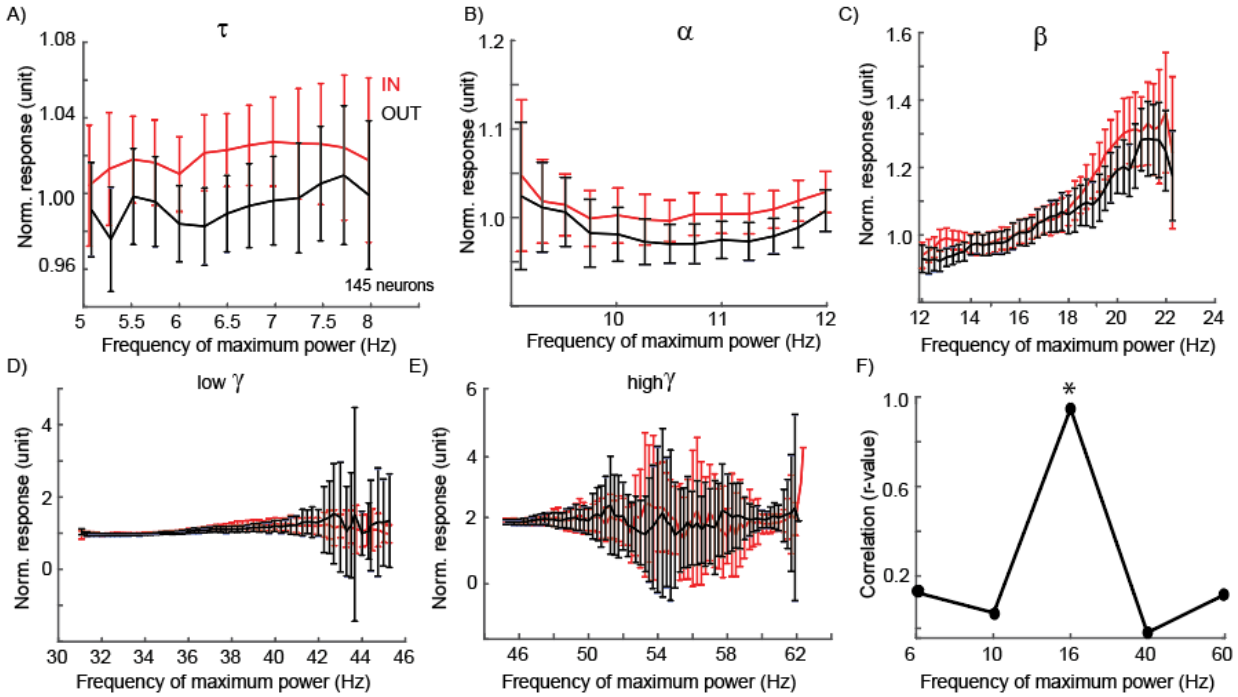


**Figure S16) Correlation between firing rate and peak frequency is specific to β frequency oscillations.** We examined multiple frequency bands to determine whether the relationship between the frequency of LFP oscillations and the firing rate of V4 neurons was general, or unique to the β band. A-E) The relationship between firing rate and the peak frequency during the delay period in different frequency bands: 4-8 Hz (theta, A), 8-12 Hz (alpha, B), 12-30 Hz (beta, C), 30-50 Hz (low gamma, D), 50-70 Hz (high gamma, E). As in figure 4G, we show average normalized response as a function of peak frequency, pooled across subsamples of trials from each of 145 V4 neurons (100 subsamples/neuron), during the IN (red) and OUT (blue) conditions. Plot shows mean±SE for all subsamples with the peak frequency indicated on the x-axis; actual subsample peak frequencies did not always span the full range of the band examined. F) The correlation between firing rate and frequency of maximum power is significant for β oscillations, but not for any other frequency bands. X-axis shows the frequency band, and y-axis is the r-value of the correlation between normalized firing rate and frequency of maximum power in that frequency band (Pearson correlation, $r_{\tau}$=0.145, $p_{\tau}$=0.620; $r_{\alpha}$=0.060, $p_{\alpha}$=0.839; $r_{\beta}$=0.921, $p_{\beta}<10^{-18}$; $r_{l\gamma}$=-0.024, $p_{l\gamma}$=0.836; $r_{h\gamma}$=0.136, $p_{h\gamma}$=0.347). These results show that the correlation between V4 LFP frequency and firing rate was unique to the β frequency range.
